# Antigenic variation of SARS-CoV-2 in response to immune pressure

**DOI:** 10.1101/2020.07.15.204610

**Authors:** Diego Forni, Rachele Cagliani, Chiara Pontremoli, Alessandra Mozzi, Uberto Pozzoli, Mario Clerici, Manuela Sironi

## Abstract

The ongoing evolution of SARS-CoV-2 is expected to be at least partially driven by the selective pressure imposed by the human immune system. We exploited the availability of a large number of high-quality SARS-CoV-2 genomes, as well as of validated epitope predictions, to show that B cell epitopes in the spike glycoprotein (S) and in the nucleocapsid protein (N) have higher diversity than non-epitope positions. Similar results were obtained for other human coronaviruses. Conversely, in the SARS-CoV-2 population, epitopes for CD4^+^ and CD8^+^ T cells were not more variable than non-epitope positions. A significant reduction in epitope variability was instead observed for some of the most immunogenic proteins (S, N, ORF8, and ORF3a). Analysis over longer evolutionary time-frames indicated that this effect is not due to differential constraints. These data indicate that SARS-CoV-2 is evolving to elude the host humoral immune response, whereas recognition by T cells might benefit the virus.

## Introduction

The COVID-19 pandemic is caused by a novel coronavirus named SARS-CoV-2 (Coronaviridae Study Group of the International Committee on Taxonomy,of Viruses, 2020). Most likely, SARS-CoV-2 originated and evolved in bats, eventually spilling over to humans, either directly or through an intermediate host (Killerby et al., 2020; Lam et al., 2020; Liu et al., 2020; Sironi et al., 2020; Wong et al., 2020; Xiao et al., 2020; Zhou et al., 2020a). Sustained human-to-human transmission determined the global spread of the virus, which has now resulted in an unprecedented global sanitary crisis. In fact, whereas the majority of COVID-19 cases are relatively mild, a significant proportion of patients develop a serious, often fatal illness, characterized by acute respiratory distress syndrome (Wu and McGoogan, 2020). Both viral-induced lung pathology and overactive immune responses are thought to contribute to disease severity (St John and Rathore, 2020; Vabret et al., 2020).

Ample evidence suggests that coronaviruses can easily cross species barriers and have high zoonotic potential. Indeed, seven coronaviruses are known to infect humans and all of them originated in animals (Cui et al., 2019; Forni et al., 2017; Ye et al., 2020). Among these, HCoV-OC43, HCoV-HKU1, HCoV-NL63 and HCoV-229E have been circulating for decades in human populations and usually cause limited disease (Bucknall et al., 1972; Forni et al., 2017; Woo et al., 2005). They are thus referred to as “common cold” coronaviruses. Conversely, MERS-CoV and SARS-CoV, whose emergence in the 2000s preceded that of SARS-CoV-2, can cause serious illness and respiratory distress syndrome in a non-negligible proportion of infected individuals (Petrosillo et al., 2020). Like all coronaviruses, these human-infecting viruses have positive-sense, single stranded RNA genomes. Two thirds of the coronavirus genome are occupied by two large overlapping open reading frames (ORF1a and ORF1b), that are translated into polyproteins. These latter are processed to generate 16 non-structural proteins (nsp1 to nsp16). The remaining portion of the genome includes ORFs for the structural proteins (spike, envelope, membrane, and nucleocapsid) and a variable number of accessory proteins (Cui et al., 2019; Forni et al., 2017).

Analysis of the bat viruses most closely related to SARS-CoV-2 indicated that, in analogy to SARS-CoV, the virus most likely required limited adaptation to gain the ability to infect and spread in our species (Boni et al., 2020; Cagliani et al., 2020). Nonetheless, since its introduction in human populations SARS-CoV-2 must have been subject to the selective pressure imposed by the human immune system. In fact, as with most other viruses, data from COVID-19, SARS, and MERS patients indicate that both B and T lymphocytes play a role in controlling infection (Channappanavar et al., 2014; St John and Rathore, 2020; Vabret et al., 2020).

Recent efforts predicted B cell and T cell epitopes in SARS-CoV-2 proteins (Grifoni et al., 2020a) and validated such predictions using sera from convalescent COVID-19 patients (Grifoni et al., 2020b). These works, as well as others (Farrera et al., 2020; Poh et al., 2020), revealed that the cell-mediated responses against SARS-CoV-2 are not restricted to the nucleocapsid (N) and spike (S) proteins, but rather target both structural and non-structural viral products. In parallel, analyses of B cell responses in SARS-CoV-2 infected patients showed that the S and N proteins are the major target of the antibody response and identified specific B cell epitopes in the S protein (Farrera et al., 2020; Jiang et al., 2020; Poh et al., 2020). We exploited this growing wealth of information to investigate whether, after a few months of sustained transmission, the selective pressure exerted by the human adaptive immune response is already detectable in the SARS-CoV-2 population and to investigate how the virus is evolving in response to such a pressure.

## Results

### Antigenic variability of SARS-CoV-2 proteins

To analyze B cell epitope diversity in SARS-CoV-2, we randomly selected 10,000 high-quality viral genomes from those available in the GISAID database (as of June 5^th^, 2020) (Elbe and Buckland-Merrett, 2017). Potential epitopes were predicted using Immune Epitope Database (IEDB) tools, as previously described (Grifoni et al., 2020a). Specifically, because they are the major targets of the humoral immune response (Channappanavar et al., 2014; St John and Rathore, 2020; Vabret et al., 2020), we predicted both linear and conformational B epitopes for the S and N proteins, whereas only linear epitopes were predicted for the other viral proteins (Table S1). A good correspondence was observed between B cell epitope predictions for the S protein and epitopes identified in two works that systematically mapped antibody responses in the sera of convalescent COVID-19 patients (Farrera et al., 2020; Poh et al., 2020) (Figure 1).

**Figure 1.**
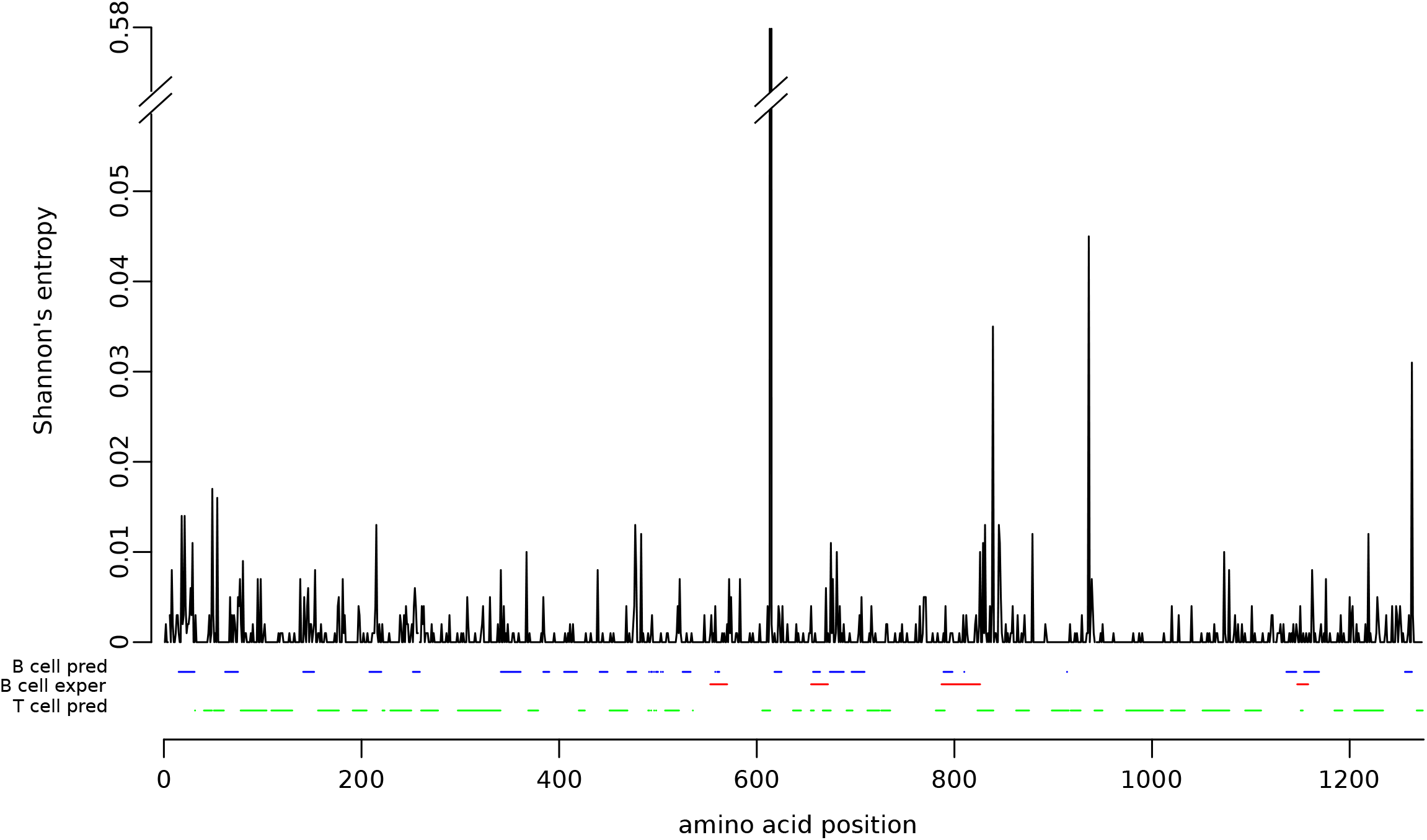
Amino acid variability of the SARS-CoV-2 spike protein. Shannon’s entropy (H) values for each amino acid position calculated using 10000 SARS-CoV-2 spike proteins are shown. B cell predicted epitopes and T cell predicted epitopes are also reported in blue and green, respectively. B cell epitopes identified in the sera of COVID-19 patients (Farrera et al., 2020; Poh et al., 2020) are also reported in red.

Variability at each amino acid site of the proteins encoded by SARS-CoV-2 was quantified using Shannon’s entropy (H). Specifically, only predicted proteins longer than 60 amino acids were analyzed. Because most positions in SARS-CoV-2 genomes are invariable across the sampled genomes, the distribution of H is zero-inflated, making the use of conventional statistical tests inappropriate (McElduff et al., 2010). We thus calculated statistical significance by permutations – i.e., by reshuffling epitope positions as amino acid stretches of the same size as the predicted epitopes. This approach also has the advantage of accounting for the possibility that, as a result of locally varying selective constraints, H is not independent among continuous protein positions.

Using this methodology, we found that, for the N and nsp16 proteins, positions mapping to predicted B cell linear epitopes are significantly more variable than those not mapping to these epitopes. Higher diversity of B cell epitopes was also observed for S, although it did not reach statistical significance (Figure 2). However, the H distribution for the spike protein includes a clear outlier represented by position 614 (Figure 1). Recent works indicated that the D614G variant, which is now prevalent worldwide, enhances viral infectivity without affecting neutralization by convalescent patient plasma (Korber et al., 2020; Yurkovetskiy et al., 2020; Zhang et al., 2020b). Hence, the frequency increase of this variant is unlikely to be related to immune evasion. We thus repeated the analyses after excluding position 614 and we observed that predicted B cell linear epitopes in the spike protein are significantly more variable than non-epitope positions (Figure 2). The same analysis for B cell conformational epitopes in the N and S proteins indicated a similar trend, although statistical significance was not reached (not shown). This is most likely due to the small number of positions in these epitopes. Overall, these data fit very well with the observation that most humoral immune responses against SARS-CoV-2 and other human coronaviruses are directed against the S and N proteins (Farrera et al., 2020; Jiang et al., 2020; Poh et al., 2020). These results also support the notion that the selective pressure exerted by the human antibody response is already detectable in the SARS-CoV-2 population. We next assessed whether epitopes for cell-mediated immune responses are also more variable than non-epitope positions. We thus retrieved predicted CD4^+^ and CD8^+^ T cell epitopes from a previous work (Grifoni et al., 2020a). These epitope predictions were shown to be reliable, as they capture a significant proportion of T cell responses in the sera of convalescent COVID-19 patients (Grifoni et al., 2020b). Analysis of entropy values indicated that CD4^+^ T cell epitopes are significantly less variable than non-epitope positions for the N and nsp16 proteins (Figure 2). A similar trend was observed for ORF8, E, and S, although significance was not reached. Reduction of variability was also observed for CD8^+^ T cell epitopes for the N protein, as well as for ORF3a. Higher variability in epitope positions was observed for nsp8 and nsp14 for CD4^+^ T cells alone (Figure 2). Because several epitopes for T cells co-map with B cell epitopes, which tend to show higher diversity, we compared positions within CD4^+^ or CD8^+^ T cell epitopes only (not overlapping with B cell epitopes) with positions not mapping to any of these epitopes. A significant reduction of variability was observed for S, N, ORF8, nsp15, and nsp16, whereas higher diversity was still evident for nsp8 (Figure 2).

**Figure 2.**
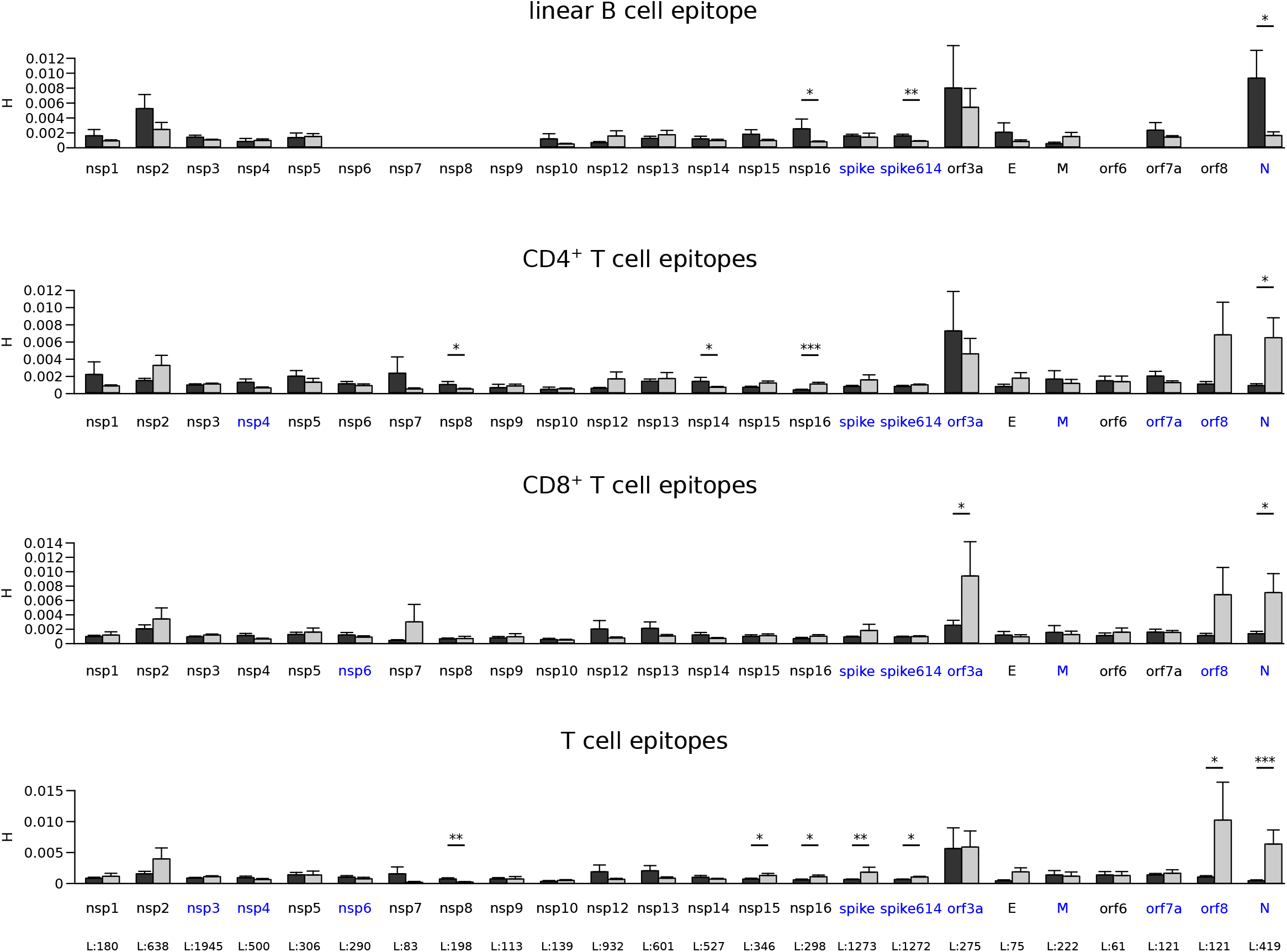
Variability of epitope and non-epitope positions among SARS-CoV-2 proteins. Shannon’s entropy (H) mean values along with standard errors are shown for all SARS-CoV-2 proteins longer than 60 residues. Epitope positions are shown in dark gray, non-epitopes in light gray. Significant comparisons, calculated by a permutation approach, are indicated with asterisks (*, P < 0.05; **, P < 0.01; ***, P < 0.001). Immunogenic proteins are shown in blue and the length of each protein is reported in the bottom panel.

Overall, these data indicate that T cell epitopes in the most immunogenic SARS-CoV-2 proteins (S, N, ORF3a, and ORF8) (Grifoni et al., 2020b; Peng et al., 2020b) tend to be more conserved than non-epitopes. However, this was not the case for other proteins targeted by T cell responses, namely M, ORF7a, nsp3, nsp4, and nsp6.

Clearly, protein sequence variability is strongly influenced by functional and structural constraints. We thus reasoned that if the observations reported above were secondary to the incidental co-localization of T cell epitopes with more constrained regions, a similar pattern should be observed for H values calculated on an alignment of proteins from other sarbecoviruses. In fact, all these viruses, with the exclusion of SARS-CoV, were sampled from bats. Thus, whereas structural/functional constraints are expected to be maintained across long evolutionary time frames, the pressure exerted by the human cell-mediated immune response is not, as, in different species, antigen processing within host cells results in the preferential presentation of diverse viral epitopes to T lymphocytes depending on the MHC gene repertoire and on distinct preferences of the antigen processing pathway (Abduriyim et al., 2019; Burgevin et al., 2008; Hammer et al., 2007; Lu, Dan AND Liu, Kefang AND Zhang, Di AND Yue, Can AND Lu, Qiong AND Cheng, Hao AND Wang, Liang AND Chai, Yan AND Qi, Jianxun AND Wang, Lin-Fa AND Gao, George F. AND Liu,William J., 2019; Wynne et al., 2016). Conversely, epitopes for antibodies tend to be conserved across species (Tse et al., 2017; Wiehe et al., 2014) and consequently the selective pressure acting on these positions is expected to be constant across time and hosts.

We thus aligned the SARS-CoV-2 reference sequences of proteins showing decreased or increased variability in T cell epitopes with those of 45 sarbecoviruses (Table S2). Calculation of H indicated a significant difference only for CD4^+^ T cell epitopes in the N protein. Conversely, B cell epitopes were more variable than non-epitope positions for the S, N and nsp16 proteins (Figure 3). Overall, these results indicate that the variability within SARS-CoV-2 T cell epitopes is not primarily driven by functional/structural constraints, but most likely results from the interaction with the human adaptive immune response.

**Figure 3.**
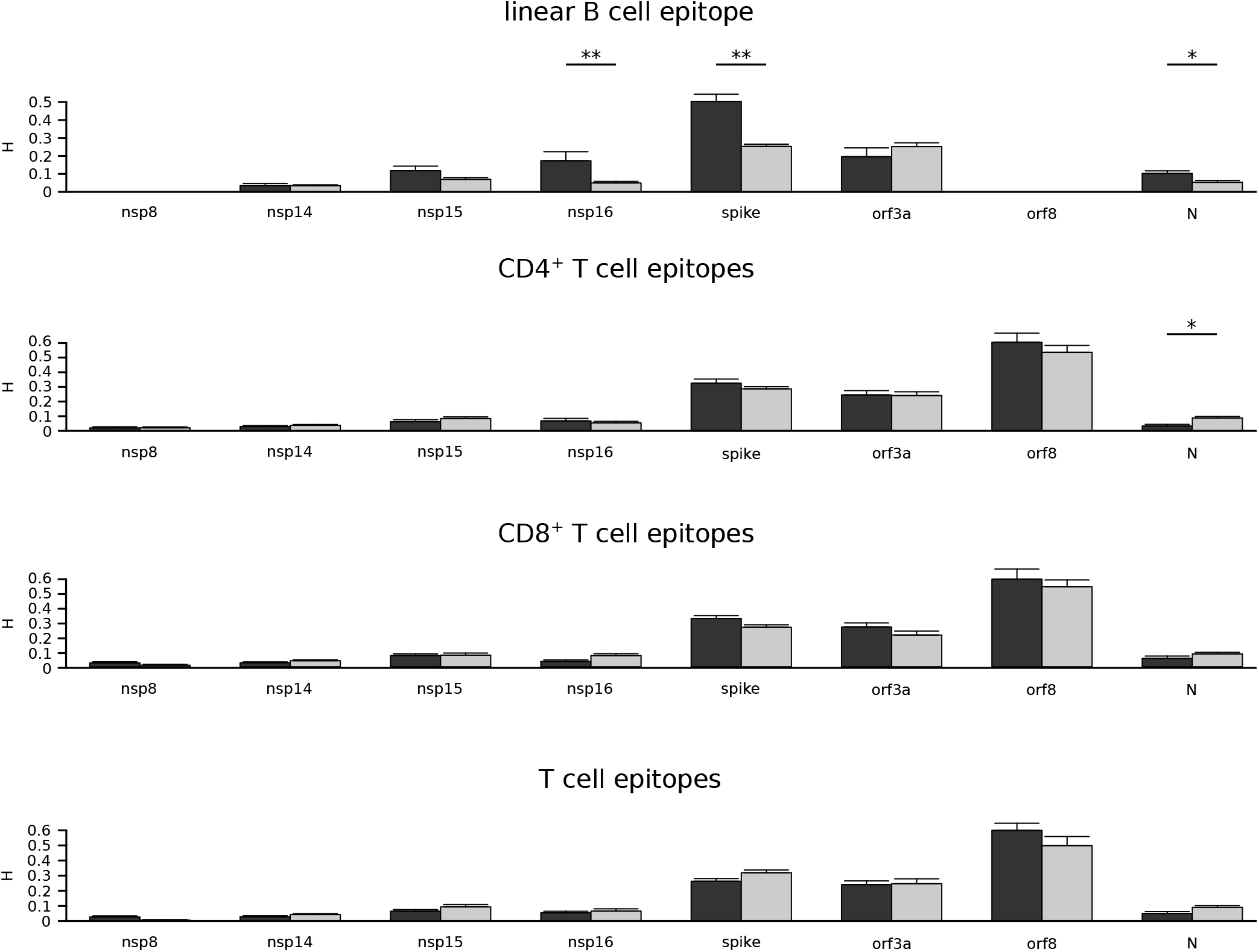
Variability of epitope and non-epitope positions among sarbecoviruses. Shannon’s entropy (H) mean values along with standard errors are shown for a set of sarbecovirus ORFs. SARS-CoV-2 epitope positions are shown in dark gray, non-epitopes in light gray. Significant comparisons, calculated by a permutation approach, are indicated with asterisks (*, P < 0.05; **, P < 0.01; ***, P < 0.001).

### Comparison with other human coronaviruses

Given the results above we set out to determine whether the other human coronaviruses show the same tendency of reduced and increased variability at T cell and B cell epitopes, respectively. For these viruses, analyses were restricted to the N and S proteins, as they are the most antigenic proteins and because the number of complete viral genomes is relatively limited (Table S3).

SARS-CoV, the human coronavirus most similar to SARS-CoV-2, caused the first human outbreak in 2002/2003 after a spill-over from palm civets, followed by human-to-human transmission chains (Shi and Wang, 2017). A second zoonotic transmission occurred in December 2003 and caused a limited number of cases (Shi and Wang, 2017; Wang et al., 2005). Viral genomes sampled during the second outbreak were not included in the analyses because their evolution occurred in the civet reservoir (Table S4). Four other human coronaviruses – HCoV-OC43, HCoV-HKU1 (members of the *Embecovirus* subgenus), HCoV-229E *(Duvinavirus* subgenus), and HCoV-NL63 *(Setracovirus* subgenus) – have been transmitting within human populations for at least 70 years (Forni et al., 2017). Thus, all available S and N sequences were included in the analyses (Table S4). Conversely, MERS-CoV displays limited ability of human-to-human transmission and outbreaks were caused by repeated spill-over events from the camel host (Cui et al., 2019). For this reason MERS-CoV was excluded from the analyses.

Quantification of sequence variability by H calculation indicated that B cell epitopes in the S protein are significantly more variable than non-epitopes for SARS-CoV, HCoV-OC43 and HCoV-HKU1 (Figure 4). Analysis of CD4^+^ and CD8^+^ T cell epitopes in these viruses indicated no increased diversity for epitope compared to non-epitope positions, with the exclusion of the S protein of SARS-CoV for CD4^+^ T cells. However, when positions within B cell epitopes were excluded from the analysis, this difference disappeared and T cell epitopes were found to be significantly less variable than nonepitopes for the spike proteins of HCoV-HKU1 and HCoV-OC43, as well as for the N protein of HCoV-229E (figure 4). Thus, the lack of antigenic diversity at T cell epitopes is a common feature of human coronaviruses, which instead tend to maintain sequence conservation of such epitopes.

**Figure 4.**
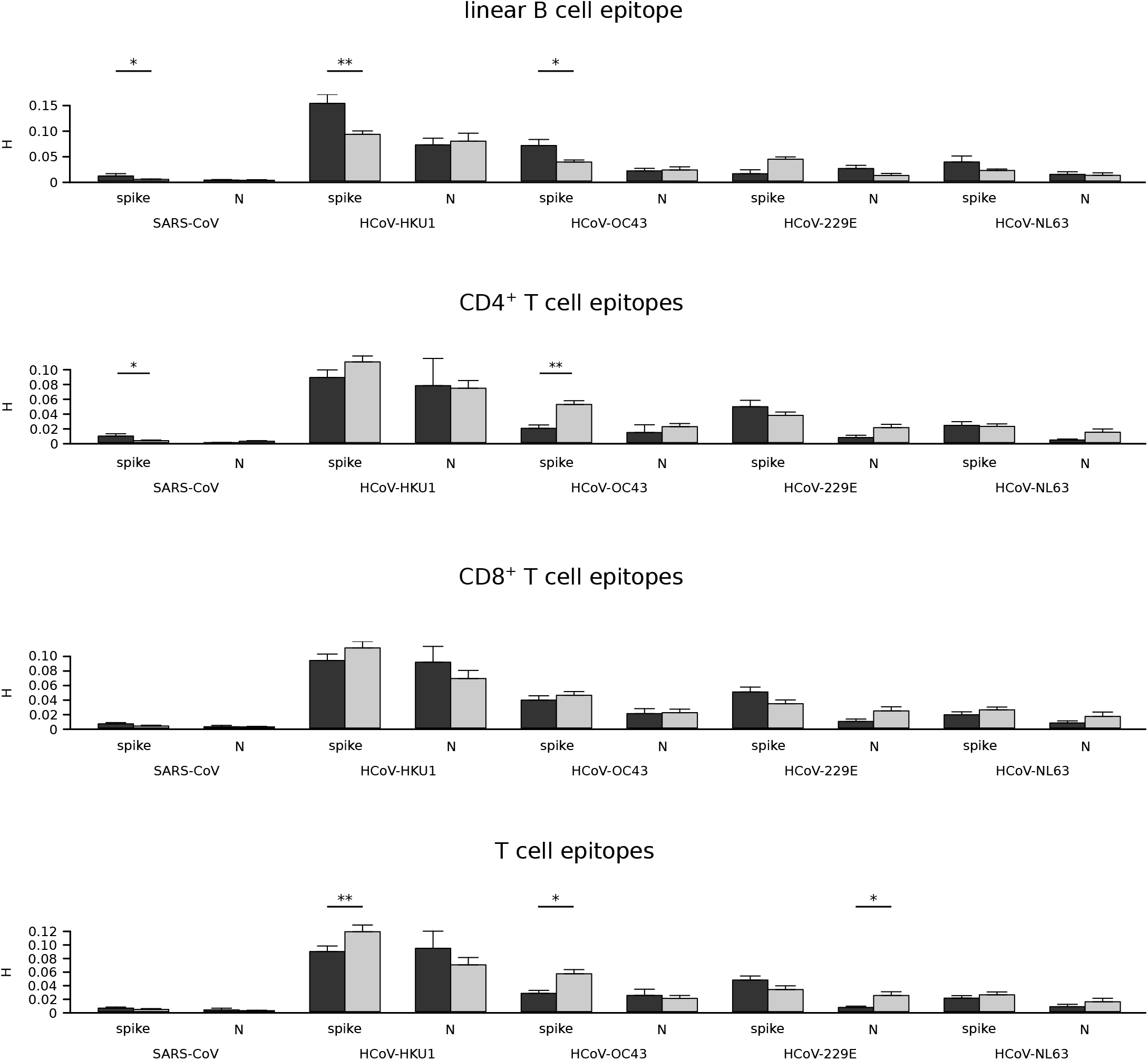
Variability of epitope and non-epitope positions among human coronaviruses. Shannon’s entropy (H) mean values along with standard errors are shown for human coronavirus spike and nucleocapsid proteins. Epitope positions are shown in dark gray, non-epitopes in light gray. Significant comparisons, calculated by a permutation approach, are indicated with asterisks (*, P < 0.05; **, P < 0.01; ***, P < 0.001).

## Discussion

The origin of SARS-CoV-2 is still uncertain and it is presently unknown whether the virus spilled over from a bat or another intermediate host. The hypothesis of a zoonotic origin is strongly supported by multiple lines of evidence, although it cannot be excluded that SARS-CoV-2 transmitted cryptically in humans before gaining the ability of spreading efficiently among people (Andersen et al., 2020; Sironi et al., 2020). Whatever the initial events associated with the early phases of the pandemic, it is clear that circulating SARS-CoV-2 viruses shared a common ancestor at the end of 2019 (Li et al., 2020; van Dorp et al., 2020). Due to its recent origin, the genetic diversity of the SARS-CoV-2 population is still limited. This is also the result of the relatively low mutation rate of coronaviruses (as compared to other RNA viruses), which encode enzymes with some proofreading ability (Denison et al., 2011; Forni et al., 2017). Nonetheless, the huge number of transmissions worldwide have allowed thousands of mutations to appear in the viral population and, thanks to enormous international sequencing efforts, more than 14,000 amino acid replacements have currently been reported (http://cov-glue.cvr.gla.ac.uk). Irrespective of the host, most variants are expected to be deleterious for viral fitness, or to have no consequences (Cagliani et al., 2020; Grubaugh et al., 2020; van Dorp et al., 2020). However, a portion of replacements may favor the virus and some of these may contribute to adaptation to the human host. In particular, the recent and ongoing evolution of SARS-CoV-2 is expected to be at least partially driven by the selective pressure imposed by the human immune system. Indeed, antigenic drift or immune evasion mutations have been reported for other zoonotic viruses such as Lassa virus (Andersen et al., 2015) and Influenza A virus (Su et al., 2015). The emergence of immune evasion variants was also observed during an outbreak of MERS-CoV in South Korea, when mutations in the spike proteins were positively selected as they facilitated viral escape from neutralizing antibodies, even though the same variants decreased binding to the cellular receptor (Kim et al., 2019; Kim et al., 2019; Kleine-Weber et al., 2019; Rockx et al., 2010). This exemplifies a phenomenon often observed in other viruses, most notably HIV-1 (Liu et al., 2007; Martinez-Picado et al., 2006; Schneidewind et al., 2007; Schneidewind et al., 2008), whereby the virus trades off immune evasion with a fitness cost. As a consequence, immune evasion mutations may be only transiently maintained in viral populations. For this reason we decided to quantify epitope variability in terms of entropy, rather than relying on measures based on substitution rates (dN/dS), which were developed for application to variants that go to fixation in different lineages over time (Kryazhimskiy and Plotkin, 2008).

The MERS-CoV mutants responsible for the outbreak in South Korea also testify the relevance of the antibody response in coronavirus control and the selective pressure imposed by humoral immunity on the virus (Kim et al., 2019; Kleine-Weber et al., 2019; Rockx et al., 2010). This is most likely the case for SARS-CoV-2, as well, as recent report indicated that the sera of most COVID-19 convalescent patients have virus-neutralization activities and antibody titers negatively correlate with viral load (OKBA et al., 2020; Vabret et al., 2020; Wu et al., 2020; Zhou et al., 2020b). Nonetheless, studies on relatively large COVID-19 patient cohorts reported that patients with severe disease display stronger IgG responses than milder cases and a negative correlation between anti-S antibody titers and lymphocyte counts was reported (Jiang et al., 2020; Vabret et al., 2020; Wu et al., 2020; Zhang et al., 2020a; Zhao et al., 2020). Consistently, asymptomatic SARS-CoV-2 infected individuals were recently reported to have lower virus-specific IgG levels than COVID-19 patients (Long et al., 2020). These observations raised concerns that humoral responses might not necessarily be protective, but rather pathogenic, either via antibody-dependent enhancement (ADE) or other mechanisms (Cao, 2020; Iwasaki and Yang, 2020; Wu et al., 2020).

Clearly, gaining insight into the dynamic interaction between SARS-CoV-2 and the human immune system is of fundamental importance not only to understand COVID-19 immunopathogenesis, but also to inform therapeutic and preventive viral control strategies. We thus exploited the availability of a large number of fully sequenced high-quality SARS-CoV-2 genomes, as well as validated predictions of B cell and T cell epitopes, to investigate whether the selective pressure exerted by the adaptive immune response is detectable in global SARS-CoV-2 population, and if the virus is evolving to evade it. Results indicated that B cell epitopes in the N and S proteins, which represent the major targets of the antibody response, have higher diversity than non-epitope positions. The same was observed for the spike proteins of HCoV-HKU1, HCoV-OC43 and SARS-CoV, although data on the latter virus should be taken with caution as they derive from a relatively small number of sequences sampled over a short time frame. Conversely, no evidence of antibody-mediated selective pressure was evident for HCoV-229E and HCoV-NL63. The reasons underlying these differences are unclear, but recent data on a relatively small population of patients with respiratory disease indicated that the titers of neutralizing antibodies against HCoV-OC43 tend to be higher compared to those against HCoV-229E and HCoV-NL63 (HCoV-HKU1 was not evaluated), suggesting the two latter viruses elicit mainly nonneutralizing responses (Gorse et al., 2020).

B cell epitopes within nsp16 were also found to be variable, although this protein was not reported to be immunogenic (Grifoni et al., 2020b). However, the antibody response to SARS-CoV-2 has presently been systematically analyzed in a relatively small number of patients and most studies focused on structural proteins. It is thus possible that, during infection, antibodies against nsp16 are raised but they have not been detected yet. An alternative possibility is that B cell epitopes in nsp16, which is highly conserved in SARS-CoV-2 strains (Cagliani et al., 2020), coincide with regions of relatively weaker constraint. This hypothesis is partially supported by the observation that these same positions also display higher diversity when entropy is calculated on an alignment of sarbecovirus nsp16 proteins. More intriguingly, this result may indicate that nsp16, together with S and N, is a target of B cell responses in the bat reservoirs. In fact, as mentioned above, antibody binding sites tend to be conserved across species (Tse et al., 2017; Wiehe et al., 2014) and thus the selective pressure exerted on B cell epitopes is likely to be constant across hosts. Whereas the immunogenicity of nsp16 remains to be evaluated, these data suggest that SARS-CoV-2 is evolving to elude the host humoral immune response. We however note that this observation does not necessarily imply that antibodies against SARS-CoV-2 are protective and it does not rule out the possibility that humoral responses contribute to COVID-19 pathogenesis.

In COVID-19 patients, antibody titers were found to correlate with the strength of virus-specific T cell responses (Ni et al., 2020). Surprisingly, we found that, in the SARS-CoV-2 population, epitopes for CD4^+^ and CD8^+^ T cells are not more variable than non-epitope positions. Conversely, a significant reduction in epitope variability was observed for a subset of viral proteins, in particular for some of the most immunogenic ones (S, N, ORF8, and ORF3a) (Grifoni et al., 2020b; Peng et al., 2020a). To check that the result was not due to stronger structural/functional constraints acting on epitope positions, we again used H values calculated on an alignment of sarbecovirus genomes, all of which, with the exclusion of SARS-CoV, were sampled in bats. T cell responses are initiated by the presentation of antigenic epitopes by MHC (major histocompatibility complex) class I and class II molecules. Different mammals have diverse MHC gene repertoires and thus present distinct antigens. In particular, recent data from various bat species indicated that many MHC class I molecules have a 3- or 5-amino acid insertion in the peptide binding pocket, resulting in very different presented peptide repertoires compared to the MHC class I molecules of other mammals (Abduriyim et al., 2019; Lu, Dan AND Liu, Kefang AND Zhang, Di AND Yue, Can AND Lu, Qiong AND Cheng, Hao AND Wang, Liang AND Chai, Yan AND Qi, Jianxun AND Wang, Lin-Fa AND Gao, George F. AND Liu,William J., 2019; Ng et al., 2016; Papenfuss et al., 2012; Wynne et al., 2016). Thus, the selective pressure acting on T cell epitopes is most likely volatile and not conserved in humans and bats. Analysis of sarbecovirus proteins indicated that, apart from CD4^+^ T cell epitopes in the N protein, the T cell epitopes predicted in SARS-CoV-2 proteins are not less diverse than non-epitope positions, suggesting that epitope conservation is not simply secondary to structural or functional constraints, but may result from interaction with human T cell responses. Of course, another possible explanation for this finding is that the prediction tools failed to identify real epitopes. However, we retrieved epitopes from a previous work and the authors validated their predictions using the sera of 20 patients who recovered from COVID-19 (Grifoni et al., 2020a; Grifoni et al., 2020b). Moreover, if a general artifact linked to epitope prediction was introduced, we would not expect to observe significant differences and not specifically in the proteins that represent the major targets of T cell responses.

Unexpected conservation of T cell epitopes was previously observed for HIV-1 and *Mycobacteriun tuberculosis* (MTB), both of which cause chronic infections in humans (Comas et al., 2010; Coscolla et al., 2015; Lindestam Arlehamn et al., 2015; Sanjuán, Rafael AND Nebot, Miguel R. AND Peris, Joan B. AND Alcamí,José, 2013). In the case of HIV-1, immune activation most likely favors the virus by increasing the rate of CD4^+^ T cell trans-infection (Sanjuán, Rafael AND Nebot, Miguel R. AND Peris, Joan B. AND Alcamí,José, 2013). Conversely, the mechanisms underlying MTB epitope conservation are not fully elucidated. A possible explanation is that conserved epitopes generate a decoy immune response and advantage the bacterium. An alternative possibility is that T cell activation results in lung tissue inflammation and damage (cavitary tuberculosis), which favors MTB transmission by aerosol (Coscolla et al., 2015; Lindestam Arlehamn et al., 2015). Although these mechanisms are unlikely to be at play in the case of SARS-CoV-2, a deregulated immune response has been associated with COVID-19 pathogenesis (Hannan et al., 2020). Specifically, recent data indicated that patients recovering from severe COVID-19 have broader and stronger T cell responses compared to mild cases (Peng et al., 2020b). This was particularly evident for responses against the S, membrane (M), ORF3a, and ORF8 proteins (Peng et al., 2020b). Although this observation might simply reflect higher viral loads in severe cases, the possibility that the T cell response itself is deleterious cannot be excluded. Moreover, the same authors reported that CD8^+^ T cells targeting different virus proteins have distinct cytokine profiles, suggesting that the virus can modulate the host immune response to its benefit (Peng et al., 2020b). Additionally, a post-mortem study on six patients who died from COVID-19 indicated that infection of macrophages can lead to activation-induced T cell death, which may eventually be responsible for lymphocytopenia (chen et al., 2020). However, we also found a trend of lower diversity of T cell epitopes for common cold coronaviruses, indicating that epitope conservation *per se* is not directly linked to disease severity. Moreover, other SARS-CoV-2 immunogenic proteins such as M and ORF7 did not show differences in T cell epitope conservation, which was instead observed for nsp16 and nsp15. These latter are not known to be T cell targets (Grifoni et al., 2020b). Clearly, further analyses will be required to clarify the significance of T cell epitope conservation in SARS-CoV-2. An interesting possibility is that both for SARS-CoV-2 and for common cold coronaviruses, conservation serves to maintain epitopes that elicit tolerizing T cell responses or induce T cells with regulatory activity. Indeed, we considered T cell epitopes as a whole, but differences exist in terms of variability and, most likely, antigenicity. This clearly represents a limitation of this study, but the modest amount of genetic diversity in the SARS-CoV-2 population does not presently allow analysis of single epitope regions. Moreover, more detailed and robust analyses will indubitably require the systematic, experimental definition of T and B cell epitopes in the SARS-CoV-2 proteome.

## Material and Methods

### Epitope Prediction

Epitope prediction was performed using different tools from The Immune Epitope Database (IEDB) (https://www.iedb.org/), as previously suggested (Grifoni et al., 2020a). Protein sequences from reference strains of human coronaviruses were used as input for all prediction analyses (SARS-CoV-2, NC_045512; SARS-CoV, NC_004718; Human coronavirus 229E, NC_002645; Human coronavirus NL63, NC_005831; Human coronavirus OC43, NC_006213; Human coronavirus HKU1, NC_006577). In particular, for linear B cell epitope prediction, we used the Bepipred Linear Epitope Prediction 2.0 tool (Jespersen et al., 2017) with a cutoff of 0.550 and epitope length > 7. Conformational B epitopes for the S and N proteins of SARS-CoV-2 were calculated using Discotope 2.0 (Kringelum et al., 2012) with a threshold = −2.5 and published 3D protein structures (PDB IDs: 6VSB, spike; 6M3M (N-term) and 7C22 (C-term), nucleocapsid protein).

SARS-CoV-2 predicted T cell epitopes were retrieved from a previous work (Grifoni et al., 2020a). For all other coronaviruses, we applied the same methodology used by Grifoni et al. (Grifoni et al., 2020a). CD4^+^ cell epitopes were predicted using TepiTool (Paul et al., 2016) with default parameters. CD8^+^ epitopes were predicted by using MHC-I Binding Predictions v2.23 tool (http://tools.iedb.org/mhci/). The NetMHCpan EL 4.0 method (Jurtz et al., 2017) was applied and the 12 most frequent HLA class I alleles in human populations (HLA-A01:01, HLA-A02:01, HLA-A03:01, HLA-A11:01, HLA-A23:01, HLA-A24:02, HLA-B07:02, HLA-B08:01, HLA-B35:01, HLA-B40:01, HLA-B44:02, HLA-B44:03) were analyzed with a 8-14 kmer range. Only epitopes with a score rank ≤ 0.1 in one of the 12 HLA classes were selected.

### Sequences and alignments

SARS-CoV-2 protein sequences were downloaded from the GISAID Initiative (https://www.gisaid.org) database (as of June, 5^th^). All protein sequences were retrieved and several filters were applied. Only complete genomes flagged as “high coverage only” and “human” were selected. Positions recommended to be masked by DeMaio and coworkers (https://virological.org/t/masking-strategies-for-sars-cov-2-alignments/480, last accessed June, 5^th^, 2020) were also removed.

Finally, for each SARS-CoV-2 protein, we selected only strains that had the same length as the protein in the SARS-CoV-2 reference strain (NC_045512), generating a set of at least 23625 sequences for each ORF. Proteins with less than 60 amino acids were excluded from the analyses.

The list of GISAID IDs along with the list of laboratories which generated the data is provided as Table S5.

For all the other human coronaviruses, as well as for a set of non-human infecting sarbecoviruses, sequences of either complete genome or single ORFs (i.e. nucleocapsid and spike protein) were retrieved from the National Center for Biotechnology Information database (NCBI, http://www.ncbi.nlm.nih.gov/). For all human coronaviruses, the only filter we applied was the host identification as “human”. SARS-CoV strains sampled during the second outbreak were excluded from the analyses. NCBI ID identifiers are listed as Table S2 and Table S4.

Alignments were generated using MAFFT (Katoh and Standley, 2013).

### Protein variability and statistical analysis

Variability at each amino acid position was estimated using the Shannon’s entropy (H) index using the Shannon Entropy-One tool from the HIV database (https://www.hiv.lanl.gov/content/index), with ambiguous character (e.g. gaps) excluded from the analysis. For SARS-CoV-2 strains, H was calculated on alignments of 10000 randomly selected sequences for each protein. For each protein we evaluated the difference D between average H values at epitope and non-epitope positions.

Most positions of analyzed viruses are invariable along the alignments, so the distribution of H is zero-inflated. We thus calculated statistical significance by permutations. For each protein, the predicted epitope intervals were collapsed to a single position while non-epitope intervals were left unchanged. After randomly shuffling this collapsed sequence it was expanded back to full length and the difference between shuffled epitope and non-epitope H values was calculated. This procedure was repeated 1000 times and the proportion of repetitions showing a difference more extreme than D was reported as p-value. An in house R script was written and is available as supplementary text S1.

## Acknowledgments

We gratefully acknowledge the authors, originating and submitting laboratories of the sequences from GISAID’s EpiCoV™ database on which this research is based. This work was supported by the Italian Ministry of Health (“Ricerca Corrente 2019-2020” to MS, “Ricerca Corrente 2018-2020” to DF)

## Author Contributions

Conceptualization, DF and MS; Formal Analysis, MS, UP, and DF; Investigation, DF, RC, CP, AM, and MS; Visualization, DF, RC; Writing –Original Draft, MS and DF Writing –Review & Editing, MS, MC, RC, UP; Funding Acquisition MS and DF; Supervision, MS and MC.

