## Supplementary table 2 for "Antigenic variation of SARS-CoV-2 in response to immune pressure"

**Table S2. List of sarbecoviruses used in the study.**

| **ID** | **STRAIN** | **YEAR** | **COUNTRY** | **HOST** |
| --- | --- | --- | --- | --- |
| **Sarbecovirus** | | | | |
| KP886808 | YNLF_31C | 2013 | China | *Rhinolophus Ferrumequinum* |
| KJ473811 | BtRf-JL2012 | 2012 | China | *Rhinolophus ferrumequinum* |
| NC_004718 | Tor2 | 2003 | Canada | *Homo sapiens* |
| KT444582 | WIV16 | 2013 | China | *Rhinolophus sinicus* |
| MN996532 | RaTG13 | 2013 | China | *Rhinolophus affinis* |
| MG772934 | bat-SL-CoVZXC21 | 2015 | China | *Rhinolophus sinicus* |
| MT084071 | MP789 | 2019 | China | *Manis javanica* |
| KY417145 | Rf4092 | 2012 | China | *Rhinilophus ferrumequinum* |
| GQ153547 | HKU3-12 | 2007 | China | *Rhinolophus sinicus* |
| KY417147 | Rs4237 | 2013 | China | *Rhinolophus sinicus* |
| KY417146 | Rs4231 | 2013 | China | *Rhinolophus sinicus* |
| DQ071615 | Rp3 | 2004 | China | *Rhinolophus sinicus* |
| KY417143 | Rs4081 | 2012 | China | *Rhinolophus sinicus* |
| KY417142 | As6526 | 2014 | China | *Aselliscus stoliczkanus* |
| KY417149 | Rs4255 | 2013 | China | *Rhinolophus sinicus* |
| KY417148 | Rs4247 | 2013 | China | *Rhinolophus sinicus* |
| KC881005 | RsSHC014 | 2011 | China | *Rhinolophus sinicus* |
| JX993988 | Cp/Yunnan2011 | 2011 | China | *Chaerephon plicata* |
| KC881006 | Rs3367 | 2012 | China | *Rhinolophus sinicus* |
| NC_045512 | Wuhan-Hu-1 | 2019 | China | *Homo sapiens* |
| DQ412042 | Rf1 | 2004 | China | *Rhinolophus ferrumequinum* |
| KY770860 | Jiyuan-84 | 2012 | China | *Rhinolophus ferrumequinum* |
| MG772933 | bat-SL-CoVZC45 | 2017 | China | *Rhinolophus sinicus* |
| KU973692 | F46 | 2012 | China | *Rhinolophus pusillus* |
| JX993987 | Rp/Shaanxi2011 | 2011 | China | *Rhinolophus pusillus* |
| MT040335 | PCoV_GX-P5L | 2017 | China | *Manis javanica* |
| KY417144 | Rs4084 | 2012 | China | *Rhinolophus sinicus* |
| KJ473816 | BtRs-YN2013 | 2013 | China | *Rhinolophus sinicus* |
| MK211378 | BtRs-BetaCoV/YN2018D | 2016 | China | *Rhinolophus affinis* |
| KJ473815 | BtRs-GX2013 | 2013 | China | *Rhinolophus sinicus* |
| DQ412043 | Rm1 | 2004 | China | *Rhinolophus macrotis* |
| GQ153539 | HKU3-4 | 2005 | China | *Rhinolophus sp.* |
| FJ588686 | Rs672 | 2006 | China | *Rhinolophus sinicus* |
| MK211376 | BtRs-BetaCoV/YN2018B | 2016 | China | *Rhinolophus affinis* |
| MK211377 | BtRs-BetaCoV/YN2018C | 2016 | China | *Rhinolophus affinis* |
| MK211374 | BtRl-BetaCoV/SC2018 | 2016 | China | *Rhinolophus sp.* |
| GQ153543 | HKU3-8 | 2006 | China | *Rhinolophus sp.* |
| KY417151 | Rs7327 | 2014 | China | *Rhinolophus sinicus* |
| KJ473814 | BtRs-HuB2013 | 2013 | China | *Rhinolophus sinicus* |
| MK211375 | BtRs-BetaCoV/YN2018A | 2016 | China | *Rhinolophus affinis* |
| KF294457 | Longquan-140 | 2012 | China | *Rhinolophus monoceros* |
| KY770859 | Anlong-112 | 2013 | China | *Rhinolophus sinicus* |
| KF569996 | LYRa11 | 2011 | China | *Rhinolophus affinis* |
| EPI_ISL_412977***** | hCoV-19/bat/Yunnan/RmYN02/2019 | 2019 | China | *Rhinolophus malayanus* |
| *****Originating lab: Shandong First Medical University & Shandong Academy of Medical Sciences; Submitting lab: Institute of Microbiology, Chinese Academy of Sciences; Authors: Weifeng Shi, Tao Hu, Hong Zhou, Juan Li, Xing Chen, Alice Catherine Hughes, Yuhai Bi | | | | |
