## Supplementary table 4 for "Antigenic variation of SARS-CoV-2 in response to immune pressure"

**Table S4. List of HCoV-229E, HCoV-NL63, SARS-CoV, HCoV-OC43, and HCoV-HKU1 strains used in the study.**

| ***Alphacoronavirus***  **Subgenus: *Duvinacovirus*** | | | | | |  |
| --- | --- | --- | --- | --- | --- | --- |
| **Human coronavirus 229E** | | | | | |  |
| **ID** | **STRAIN** | | **YEAR** | | **COUNTRY** | **ORFs** |
| NC_002645 | 229E | | NA | | NA | Spike, N |
| KY369909 | HCoV_229E/Seattle/USA/SC677/2016 | | 2016 | | USA | Spike, N |
| KF293666 | 229E K22-p11 | | NA | | Sweden | Spike, N |
| KF293665 | 229E clone p11 | | NA | | Sweden | Spike, N |
| KF293664 | 229E clone p0 | | NA | | Sweden | Spike, N |
| KU291448 | HCoV-229E/BN1/GER/2015 | | 2015 | | Germany | Spike, N |
| KY674914 | HCoV_229E/Seattle/USA/SC399/2016 | | 2016 | | USA | Spike, N |
| MT438699 | HCoV-229E/USA/ACRI_0246/2017 | | 2017 | | USA | Spike, N |
| MN369046 | HCoV_229E/Seattle/USA/SC9724/2018 | | 2018 | | USA | Spike, N |
| KF514431 | 229E/human/USA/953-49/1995 | | 1995 | | USA | Spike, N |
| KY674919 | N08-434B | | 2016 | | USA | Spike, N |
| KF514432 | 229E/human/USA/932-72/1993 | | 1993 | | USA | Spike, N |
| KY621348 | HCoV_229E/Seattle/USA/SC379/2016 | | 2016 | | USA | Spike, N |
| KF514429 | 229E/human/USA/892-11/1989 | | 1989 | | USA | Spike, N |
| KF514433 | 229E/human/USA/933-40/1993 | | 1993 | | USA | Spike, N |
| KF514430 | 229E/human/USA/933-50/1993 | | 1993 | | USA | Spike, N |
| KY369910 | HCoV_229E/Seattle/USA/SC1143/2016 | | 2016 | | USA | Spike, N |
| KY369908 | HCoV_229E/Seattle/USA/SC579/2016 | | 2016 | | USA | Spike, N |
| KY369912 | HCoV_229E/Seattle/USA/SC9731/2016 | | 2016 | | USA | Spike, N |
| JX503061 | J0304 | | 2009 | | Italy | Spike, N |
| KY369911 | HCoV_229E/Seattle/USA/SC1212/2016 | | 2016 | | USA | Spike, N |
| KY369914 | HCoV_229E/Seattle/USA/SC9773/2016 | | 2016 | | USA | Spike, N |
| KY983587 | HCoV_229E/Seattle/USA/SC3112/2015 | | 2015 | | USA | Spike, N |
| KY369913 | HCoV_229E/Seattle/USA/SC1073/2016 | | 2016 | | USA | Spike, N |
| KY967357 | HCoV_229E/Seattle/USA/SC2872/2015 | | 2015 | | USA | Spike, N |
| KY684760 | HCoV_229E/Seattle/USA/SC2282/2017 | | 2016 | | USA | Spike, N |
| JX503060 | 0349 | | 2010 | | Netherlands | Spike, N |
| MN306046 | HCoV_229E/Seattle/USA/SC0865/2019 | | 2019 | | USA | Spike, N |
| KY996417 | 229E/UF-1/2016 | | 2016 | | USA | Spike, N |
| MF542265 | 229E/Haiti-1/2016 | | 2016 | | Haiti | Spike, N |
| MT438698 | HCoV-229E/USA/ACRI_0256/2017 | | 2017 | | USA | Spike, N |
| MT438696 | HCoV-229E/USA/UNM_0186/2016 | | 2016 | | USA | Spike, N |
| MT438697 | HCoV-229E/USA/ACRI_0244/2017 | | 2017 | | USA | Spike, N |
| MT438700 | HCoV-229E/USA/UAMS_DID_0108/2017 | | 2017 | | USA | Spike, N |
| AF344186 | RW Stock | | NA | | NA | Spike |
| AF344187 | P11A | | NA | | NA | Spike |
| AF344188 | P11B | | NA | | NA | Spike |
| AF344189 | P100E | | NA | | NA | Spike |
| AB691763 | ATCC VR-740 | | NA | | NA | Spike |
| MH048989 | France/Lille/2014 | | 2014 | | France | Spike |
| DQ243964 | HCoV-229E-11/6/79 | | NA | | Australia | Spike |
| DQ243965 | HCoV-229E-16/6/82 | | NA | | Australia | Spike |
| DQ243966 | HCoV-229E-22/9/82 | | NA | | Australia | Spike |
| DQ243967 | HCoV-229E-6/10/82 | | NA | | Australia | Spike |
| DQ243968 | HCoV-229E-21/10/82 | | NA | | Australia | Spike |
| DQ243969 | HCoV-229E-8/11/82a | | NA | | Australia | Spike |
| DQ243970 | HCoV-229E-8/11/82b | | NA | | Australia | Spike |
| DQ243971 | HCoV-229E-29/7/84 | | NA | | Australia | Spike |
| DQ243972 | HCoV-229E-5/9/84 | | NA | | Australia | Spike |
| DQ243973 | HCoV-229E-24/6/90 | | NA | | Australia | Spike |
| DQ243974 | HCoV-229-25/6/92 | | NA | | Australia | Spike |
| DQ243975 | HCoV-229E-12/5/92 | | NA | | Australia | Spike |
| DQ243976 | HCoV-229E-17/6/92 | | NA | | Australia | Spike |
| DQ243977 | HCoV-229E-8/8/01 | | NA | | Australia | Spike |
| DQ243978 | HCoV-229E-27/8/01 | | NA | | Australia | Spike |
| DQ243979 | HCoV-229E-21/6/02 | | NA | | Australia | Spike |
| DQ243980 | HCoV-229E-6/1/03 | | NA | | Australia | Spike |
| DQ243983 | HCoV-229E-30/7/03 | | NA | | Australia | Spike |
| DQ243984 | HCoV-229E-14/8/03 | | NA | | Australia | Spike |
| DQ243985 | HCoV-229E-19/8/03 | | NA | | Australia | Spike |
| DQ243986 | HCoV-229E-25/8/03 | | NA | | Australia | Spike |
| AB691764 | Sendai-H/1121/04 | | 2004 | | Japan | Spike |
| AB691765 | Sendai-H/826/04 | | 2004 | | Japan | Spike |
| AB691766 | Sendai-H/1948/04 | | 2004 | | Japan | Spike |
| AB691767 | Niigata/01/08 | | 2008 | | Japan | Spike |
| DQ243963 | ATCC VR-740 | | NA | | NA | Spike |
| KM055531 | 11949_2011 | | 2011 | | China | Spike |
| KM055532 | 1512A_2009 | | 2009 | | China | Spike |
| KM055533 | 1518A_2009 | | 2009 | | China | Spike |
| KM055534 | 1546A_2009 | | 2009 | | China | Spike |
| KM055535 | 1649A_2009 | | 2009 | | China | Spike |
| KM055536 | 1658A_2009 | | 2009 | | China | Spike |
| KM055537 | 1667A_2009 | | 2009 | | China | Spike |
| KM055538 | 1761A_2010 | | 2010 | | China | Spike |
| KM055539 | 1781A_2010 | | 2010 | | China | Spike |
| KM055540 | 1782A_2010 | | 2010 | | China | Spike |
| KM055541 | 1827A_2010 | | 2010 | | China | Spike |
| KM055542 | 1829A_2010 | | 2010 | | China | Spike |
| KM055543 | 1990A_2010 | | 2010 | | China | Spike |
| KM055544 | 2764A_2011 | | 2011 | | China | Spike |
| KM055545 | 8339_2009 | | 2009 | | China | Spike |
| KM055546 | 8373_2009 | | 2009 | | China | Spike |
| KM055547 | 8425_2009 | | 2009 | | China | Spike |
| KM055548 | 2861A_2011 | | 2011 | | China | Spike |
| KM055549 | 2961A_2011 | | 2011 | | China | Spike |
| KM055550 | 2981A_2011 | | 2011 | | China | Spike |
| KM055551 | 359A_2007 | | 2007 | | China | Spike |
| KM055552 | 423A_2007 | | 2007 | | China | Spike |
| KM055553 | 1482A_2009 | | 2009 | | China | Spike |
| KM055554 | 494A_2008 | | 2008 | | China | Spike |
| KM055555 | 507A_2008 | | 2008 | | China | Spike |
| KM055556 | 693A_2008 | | 2008 | | China | Spike |
| KM055560 | 1424A_2009 | | 2009 | | China | Spike |
| KM055559 | 857_2005 | | 2005 | | China | Spike |
| KM055558 | 8348_2009 | | 2009 | | China | Spike |
| KM055557 | 748_2005 | | 2005 | | China | Spike |
| KT359752 | 12MYKL0011 | | 2012 | | Malaysia | N |
| KT359753 | 12MYKL0051 | | 2012 | | Malaysia | N |
| KT359754 | 12MYKL0052 | | 2012 | | Malaysia | N |
| KT359756 | 12MYKL0126 | | 2012 | | Malaysia | N |
| KT359757 | 12MYKL0335 | | 2012 | | Malaysia | N |
| KT359758 | 12MYKL0367 | | 2012 | | Malaysia | N |
| KT359759 | 12MYKL0415 | | 2012 | | Malaysia | N |
| KT359760 | 12MYKL0543 | | 2012 | | Malaysia | N |
| KT359761 | 12MYKL0667 | | 2012 | | Malaysia | N |
| KT359763 | 12MYKL1102 | | 2012 | | Malaysia | N |
| KT359765 | 12MYKL1586 | | 2012 | | Malaysia | N |
| KT359764 | 12MYKL1271 | | 2012 | | Malaysia | N |
| KT359766 | 12MYKL1616 | | 2012 | | Malaysia | N |
| KT359767 | 13MYKL1719 | | 2013 | | Malaysia | N |
| DQ243939 | ATCC VR-740 | | NA | | NA | N |
| DQ243940 | HCoV-229E-11/6/79 | | NA | | Australia | N |
| DQ243942 | HCoV-229E-16/6/82 | | NA | | Australia | N |
| DQ243943 | HCoV-229E-6/10/82 | | NA | | Australia | N |
| DQ243944 | HCoV-229E-21/10/82 | | NA | | Australia | N |
| DQ243945 | HCoV-229E-8/11/82a | | NA | | Australia | N |
| DQ243946 | HCoV-229E-8/11/82b | | NA | | Australia | N |
| DQ243947 | HCoV-229E-5/9/84 | | NA | | Australia | N |
| DQ243948 | HCoV-229E-29/7/84 | | NA | | Australia | N |
| DQ243949 | HCoV-229E-24/6/90 | | NA | | Australia | N |
| DQ243950 | HCoV-229E-12/5/92 | | NA | | Australia | N |
| DQ243951 | HCoV-229E-17/6/92 | | NA | | Australia | N |
| DQ243952 | HCoV-229E-25/6/92 | | NA | | Australia | N |
| DQ243954 | HCoV-229E-27/8/01 | | NA | | Australia | N |
| DQ243955 | HCoV-229E-6/1/03 | | NA | | Australia | N |
| DQ243956 | HCoV-229E-24/4/03 | | NA | | Australia | N |
| DQ243957 | HCoV-229E-30/7/03 | | NA | | Australia | N |
| DQ243958 | HCoV-229E-19/8/03 | | NA | | Australia | N |
| DQ243959 | HCoV-229E-14/8/03 | | NA | | Australia | N |
| DQ243960 | HCoV-229E-28/2/03 | | NA | | Australia | N |
| DQ243961 | HCoV-229E-25/8/03 | | NA | | Australia | N |
| DQ243962 | HCoV-229E-20/1/04 | | NA | | Australia | N |
| KM055501 | 1512A_2009 | | 2009 | | China | N |
| KM055502 | 1518A_2009 | | 2009 | | China | N |
| KM055503 | 1546A_2009 | | 2009 | | China | N |
| KM055504 | 1658A_2009 | | 2009 | | China | N |
| KM055505 | 1667A_2009 | | 2009 | | China | N |
| KM055506 | 1761A_2010 | | 2010 | | China | N |
| KM055507 | 1781A_2010 | | 2010 | | China | N |
| LC005740 | 882-Yamagata-2010 | | 2010 | | Japan | N |
| LC005741 | 1060-Yamagata-2010 | | 2010 | | Japan | N |
| LC005739 | 777-Yamagata-2010 | | 2010 | | Japan | N |
| LC005738 | 775-Yamagata-2010 | | 2010 | | Japan | N |
| LC005737 | 756-Yamagata-2010 | | 2010 | | Japan | N |
| LC005736 | 452-Yamagata-2008 | | 2008 | | Japan | N |
| KT359769 | 13MYKL1883 | | 2013 | | Malaysia | N |
| DQ243941 | HCoV-229E-22/9/82 | | NA | | Australia | N |
| KT359768 | 13MYKL1847 | | 2013 | | Malaysia | N |
| KT359762 | 12MYKL1076 | | 2012 | | Malaysia | N |
| KT359755 | 12MYKL0106 | | 2012 | | Malaysia | N |
| KM055530 | 8373_2009 | | 2009 | | China | N |
| KM055529 | 2981A_2011 | | 2011 | | China | N |
| KM055528 | 2861A_2011 | | 2011 | | China | N |
| KM055527 | 1649A_2009 | | 2009 | | China | N |
| KM055526 | 1482A_2009 | | 2009 | | China | N |
| KM055525 | 1424A_2009 | | 2009 | | China | N |
| KM055524 | 11949_2011 | | 2011 | | China | N |
| KM055523 | 857_2005 | | 2005 | | China | N |
| KM055522 | 8425_2009 | | 2009 | | China | N |
| KM055521 | 8348_2009 | | 2009 | | China | N |
| KM055520 | 8339_2009 | | 2009 | | China | N |
| KM055519 | 748_2005 | | 2005 | | China | N |
| KM055518 | 693A_2008 | | 2008 | | China | N |
| KM055517 | 507A_2008 | | 2008 | | China | N |
| KM055516 | 494A_2008 | | 2008 | | China | N |
| KM055515 | 432A_2007 | | 2007 | | China | N |
| KM055514 | 359A_2007 | | 2007 | | China | N |
| KM055513 | 2961A_2011 | | 2011 | | China | N |
| KM055512 | 2764A_2011 | | 2011 | | China | N |
| KM055511 | 1990A_2010 | | 2010 | | China | N |
| KM055510 | 1829A_2010 | | 2010 | | China | N |
| KM055509 | 1827A_2010 | | 2010 | | China | N |
| KM055508 | 1782A_2010 | | 2010 | | China | N |
| **Subgenus: *Setracovirus*** | | | | | |  |
| **Human coronavirus NL63** | | | | | |  |
| **ID** | **STRAIN** | | **YEAR** | | **COUNTRY** | **ORFs** |
| NC_005831 | Amsterdam I | | NA | | NA | Spike, N |
| MG428704 | Kilifi_HH_5402_20-May-2010 | | 2010 | | Kenya | Spike, N |
| MG428702 | Kilifi_HH_3807_11-May-2010 | | 2010 | | Kenya | Spike, N |
| MG428703 | Kilifi_HH_0511_01-Jun-2010 | | 2010 | | Kenya | Spike, N |
| MG428706 | Kilifi_HH_3808_24-May-2010 | | 2010 | | Kenya | Spike, N |
| MG428700 | Kilifi_HH_1602_01-Jun-2010 | | 2010 | | Kenya | Spike, N |
| MG428701 | Kilifi_HH_0512_04-Jun-2010 | | 2010 | | Kenya | Spike, N |
| MG428705 | Kilifi_HH_0522_21-May-2010 | | 2010 | | Kenya | Spike, N |
| AY518894 | Human group 1 coronavirus associated with pnuemonia | | NA | | Netherlands | Spike, N |
| CS124012 | Sequence 55 from Patent EP1553169 | | NA | | NA | Spike, N |
| DJ009246 | NA | | NA | | NA | Spike, N |
| JX504050 | NL63/RPTEC/2004 | | 2004 | | USA | Spike, N |
| KT266906 | HCoV NL63/Haiti-1/2015 | | 2015 | | Haiti | Spike, N |
| KT381875 | NL63/UF-1/2015 | | 2015 | | USA | Spike, N |
| KU521535 | NL63/UF-2/2015 | | 2015 | | USA | Spike, N |
| KX179500 | NL63/UF-2/2015 | | 2015 | | USA | Spike, N |
| DQ445911 | Amsterdam 057 | | NA | | Netherlands | Spike, N |
| KY554971 | N07-468B_176X | | 2016 | | USA | Spike, N |
| MK334045 | ChinaGD05 | | 2018 | | China | Spike, N |
| KY554970 | N07-324B_182X | | 2016 | | USA | Spike, N |
| JX104161 | CBJ 037 | | 2008 | | China | Spike, N |
| JX524171 | CBJ123 | | 2009 | | China | Spike, N |
| MN306040 | HCoV_NL63/Seattle/USA/SC0768/2019 | | 2019 | | USA | Spike, N |
| DQ445912 | Amsterdam 496 | | NA | | Netherlands | Spike, N |
| MG772808 | CN0601/14 | | 2014 | | South Korea | Spike, N |
| MK334047 | ChinaGD04 | | 2018 | | China | Spike, N |
| MK334046 | ChinaGD01 | | 2018 | | China | Spike, N |
| KY554967 | N06-1144B | | 2016 | | USA | Spike, N |
| KY554968 | N07-185B | | 2016 | | USA | Spike, N |
| MG428707 | Kilifi_HH_5401_20-May-2010 | | 2010 | | Kenya | Spike, N |
| MK334043 | ChinaGD02 | | 2018 | | China | Spike, N |
| MK334044 | ChinaGD03 | | 2018 | | China | Spike, N |
| KY829118 | N07-262B | | 2015 | | USA | Spike, N |
| MG428699 | Kilifi_HH_5709_19-May-2010 | | 2010 | | Kenya | Spike, N |
| KY554969 | N07-196B | | 2016 | | USA | Spike, N |
| JQ900259 | NL63/DEN/2005/449 | | 2005 | | USA | Spike, N |
| KY674916 | N07-64B | | 2016 | | USA | Spike, N |
| KY983586 | HCoV_NL63/Seattle/USA/SC2940/2015 | | 2015 | | USA | Spike, N |
| JQ765566 | NL63/DEN/2008/16 | | 2008 | | USA | Spike, N |
| JQ765567 | NL63/DEN/2009/20 | | 2009 | | USA | Spike, N |
| JQ765568 | NL63/DEN/2005/193 | | 2005 | | USA | Spike, N |
| JQ765571 | NL63/DEN/2005/271 | | 2005 | | USA | Spike, N |
| JQ765572 | NL63/DEN/2005/347 | | 2005 | | USA | Spike, N |
| JQ765573 | NL63/DEN/2005/1062 | | 2005 | | USA | Spike, N |
| JQ765574 | NL63/DEN/2005/1862 | | 2005 | | USA | Spike, N |
| JQ765575 | NL63/DEN/2005/1876 | | 2005 | | USA | Spike, N |
| MN306018 | HCoV_NL63/Seattle/USA/SC0179/2018 | | 2018 | | USA | Spike, N |
| JQ765563 | NL63/DEN/2009/9 | | 2009 | | USA | Spike, N |
| JQ765564 | NL63/DEN/2009/14 | | 2009 | | USA | Spike, N |
| JQ765565 | NL63/DEN/2009/15 | | 2009 | | USA | Spike, N |
| JQ765569 | NL63/DEN/2005/232 | | 2005 | | USA | Spike, N |
| JQ765570 | NL63/DEN/2005/235 | | 2005 | | USA | Spike, N |
| JQ900257 | NL63/DEN/2009/31 | | 2009 | | USA | Spike, N |
| KF530105 | NL63/human/USA/012-31/2001 | | 2001 | | USA | Spike, N |
| KF530110 | NL63/human/USA/838-9/1983 | | 1983 | | USA | Spike, N |
| KF530104 | NL63/human/USA/904-20/1990 | | 1990 | | USA | Spike, N |
| KF530107 | NL63/human/USA/911-56/1991 | | 1991 | | USA | Spike, N |
| KF530108 | NL63/human/USA/891-6/1989 | | 1989 | | USA | Spike, N |
| KF530109 | NL63/human/USA/903-28/1990 | | 1990 | | USA | Spike, N |
| KF530111 | NL63/human/USA/901-24/1990 | | 1990 | | USA | Spike, N |
| KF530113 | NL63/human/USA/905-25/1990 | | 1990 | | USA | Spike, N |
| KF530114 | NL63/human/USA/891-4/1989 | | 1989 | | USA | Spike, N |
| KF530106 | NL63/human/USA/8712-17/1987 | | 1987 | | USA | Spike, N |
| KF530112 | NL63/human/USA/0111-25/2001 | | 2001 | | USA | Spike, N |
| KY674915 | N07-6B | | 2016 | | USA | Spike, N |
| MN026166 | NL63_KLF_01_2018 | | 2018 | | Kenya | Spike, N |
| JQ771055 | NL63/DEN/2010/20 | | 2010 | | USA | Spike |
| JQ771057 | NL63/DEN/2010/28 | | 2010 | | USA | Spike |
| KM055637 | 945A_2008 | | 2008 | | China | Spike |
| KM055633 | 2093A_2010 | | 2010 | | China | Spike |
| KM055647 | 188A_2007 | | 2007 | | China | Spike |
| KM055642 | 277A_2007 | | 2007 | | China | Spike |
| KM055634 | 1023A_2008 | | 2008 | | China | Spike |
| KM055636 | 940A_2008 | | 2008 | | China | Spike |
| KM055650 | 930A_2008 | | 2008 | | China | Spike |
| KM055649 | 996A_2008 | | 2008 | | China | Spike |
| JQ771059 | NL63/DEN/2010/35 | | 2010 | | USA | Spike |
| LC488388 | HCoVNL63/Tokyo/SGH-24/2018 | | 2018 | | Japan | Spike |
| KM055644 | 374A_2007 | | 2007 | | China | Spike |
| KM055640 | 225A_2007 | | 2007 | | China | Spike |
| KM055646 | 990A_2008 | | 2008 | | China | Spike |
| KM055632 | 1014A_2008 | | 2008 | | China | Spike |
| JQ771058 | NL63/DEN/2010/31 | | 2010 | | USA | Spike |
| KX179495 | NL63/Haiti-3/2015 | | 2015 | | Haiti | Spike |
| KM055635 | 1093A_2008 | | 2008 | | China | Spike |
| KM055648 | 954A_2008 | | 2008 | | China | Spike |
| KM055645 | 909A_2008 | | 2008 | | China | Spike |
| JQ771060 | NL63/DEN/2010/36 | | 2010 | | USA | Spike |
| KM055639 | 197A_2007 | | 2007 | | China | Spike |
| KX179494 | NL63/Haiti-2/2015 | | 2015 | | Haiti | Spike |
| LC488390 | HCoVNL63/Tokyo/SGH-15/2017 | | 2017 | | Japan | Spike |
| LC488389 | HCoVNL63/Tokyo/SGH-18/2018 | | 2018 | | Japan | Spike |
| KM055641 | 236A_2007 | | 2007 | | China | Spike |
| KM055638 | 180A_2007 | | 2007 | | China | Spike |
| KX179496 | NL63/Haiti-4/2015 | | 2015 | | Haiti | Spike |
| KM055643 | 971A_2008 | | 2008 | | China | Spike |
| JQ771056 | NL63/DEN/2010/25 | | 2010 | | USA | Spike |
| JQ900255 | NL63/DEN/2009/6 | | 2009 | | USA | N |
| JQ900260 | NL63/DEN/2005/1120 | | 2005 | | USA | N |
| JQ900256 | NL63/DEN/2009/22 | | 2009 | | USA | N |
| JQ900258 | NL63/DEN/2005/291 | | 2005 | | USA | N |
| KM055594 | 225A_2007 | | 2007 | | China | N |
| KX179498 | NL63/Haiti-3/2015 | | 2015 | | Haiti | N |
| KT359860 | 12MYKL1229 | | 2012 | | Malaysia | N |
| KM055607 | 1014A_2008 | | 2008 | | China | N |
| KM055595 | 236A_2007 | | 2007 | | China | N |
| KT359856 | 12MYKL1089 | | 2012 | | Malaysia | N |
| KY862066 | NL63/FRA-EPI/Caen/2010/09 | | 2010 | | France | N |
| KT359843 | 12MYKL0865 | | 2012 | | Malaysia | N |
| KT359849 | 12MYKL1002 | | 2012 | | Malaysia | N |
| KT359841 | 12MYKL0862 | | 2012 | | Malaysia | N |
| KT359853 | 12MYKL1064 | | 2012 | | Malaysia | N |
| KM055610 | 2093A_2010 | | 2010 | | China | N |
| KY862069 | NL63/FRA-EPI/Caen/2008/06 | | 2008 | | France | N |
| KM055608 | 1023A_2008 | | 2008 | | China | N |
| KT359847 | 12MYKL0964 | | 2012 | | Malaysia | N |
| KY862065 | NL63/FRA-EPI/Caen/2010/10 | | 2010 | | France | N |
| KT359851 | 12MYKL1013 | | 2012 | | Malaysia | N |
| KT359863 | 12MYKL1288 | | 2012 | | Malaysia | N |
| KT359829 | 12MYKL0280 | | 2012 | | Malaysia | N |
| KM055601 | 945A_2008 | | 2008 | | China | N |
| KM055603 | 971A_2008 | | 2008 | | China | N |
| KT359866 | 12MYKL1350 | | 2012 | | Malaysia | N |
| KM055593 | 197A_2007 | | 2007 | | China | N |
| KY862064 | NL63/FRA-EPI/Caen/2011/11 | | 2011 | | France | N |
| KT359836 | 12MYKL0779 | | 2012 | | Malaysia | N |
| KY862071 | NL63/FRA-EPI/Caen/2005/04 | | 2005 | | France | N |
| KT359832 | 12MYKL0719 | | 2012 | | Malaysia | N |
| DQ846901 | BJ8081 | | NA | | China | N |
| KY862055 | NL63/FRA-EPI/Caen/2015/20 | | 2015 | | France | N |
| KY862057 | NL63/FRA-EPI/Caen/2014/18 | | 2014 | | France | N |
| KY862060 | NL63/FRA-EPI/Caen/2013/15 | | 2013 | | France | N |
| KT359861 | 12MYKL1238 | | 2012 | | Malaysia | N |
| KY862059 | NL63/FRA-EPI/Caen/2013/16 | | 2013 | | France | N |
| KY862068 | NL63/FRA-EPI/Caen/2009/07 | | 2009 | | France | N |
| KT359869 | 13MYKL1797 | | 2013 | | Malaysia | N |
| KX179497 | NL63/Haiti-2/2015 | | 2015 | | Haiti | N |
| KT359867 | 12MYKL1528 | | 2012 | | Malaysia | N |
| KM055606 | 997A_2008 | | 2008 | | China | N |
| KT359839 | 12MYKL0809 | | 2012 | | Malaysia | N |
| KT359846 | 12MYKL0943 | | 2012 | | Malaysia | N |
| KT359840 | 12MYKL0813 | | 2012 | | Malaysia | N |
| KT359871 | 13MYKL2005 | | 2013 | | Malaysia | N |
| KY862063 | NL63/FRA-EPI/Caen/2011/12 | | 2011 | | France | N |
| KT359868 | 13MYKL1791 | | 2013 | | Malaysia | N |
| KT359833 | 12MYKL0730 | | 2012 | | Malaysia | N |
| KY862074 | NL63/FRA-EPI/Caen/2004/01 | | 2004 | | France | N |
| KT359862 | 12MYKL1253 | | 2012 | | Malaysia | N |
| KM055609 | 1093A_2008 | | 2008 | | China | N |
| KY862070 | NL63/FRA-EPI/Caen/2008/05 | | 2008 | | France | N |
| KX179499 | NL63/Haiti-4/2015 | | 2015 | | Haiti | N |
| KT359830 | 12MYKL0482 | | 2012 | | Malaysia | N |
| KY862073 | NL63/FRA-EPI/Caen/2004/02 | | 2004 | | France | N |
| KM055591 | 180A_2007 | | 2007 | | China | N |
| KT359842 | 12MYKL0863 | | 2012 | | Malaysia | N |
| EF081296 | 191 | | NA | | USA | N |
| KT359837 | 12MYKL0780 | | 2012 | | Malaysia | N |
| KM055602 | 954A_2008 | | 2008 | | China | N |
| KT359854 | 12MYKL1079 | | 2012 | | Malaysia | N |
| KT359850 | 12MYKL1007 | | 2012 | | Malaysia | N |
| KY862058 | NL63/FRA-EPI/Caen/2014/17 | | 2014 | | France | N |
| KM055611 | GS05_2009 | | 2009 | | China | N |
| KT359859 | 12MYKL1197 | | 2012 | | Malaysia | N |
| KM055599 | 930A_2008 | | 2008 | | China | N |
| KY862061 | NL63/FRA-EPI/Caen/2012/14 | | 2012 | | France | N |
| KM055598 | 909A_2008 | | 2008 | | China | N |
| KY862072 | NL63/FRA-EPI/Caen/2005/03 | | 2005 | | France | N |
| KM055605 | 996A_2008 | | 2008 | | China | N |
| KT359844 | 12MYKL0880 | | 2012 | | Malaysia | N |
| KT359870 | 13MYKL1870 | | 2013 | | Malaysia | N |
| KY862067 | NL63/FRA-EPI/Caen/2009/08 | | 2009 | | France | N |
| KT359838 | 12MYKL0802 | | 2012 | | Malaysia | N |
| KT359855 | 12MYKL1088 | | 2012 | | Malaysia | N |
| KM055597 | 374A_2007 | | 2007 | | China | N |
| KM055604 | 990A_2008 | | 2008 | | China | N |
| KT359845 | 12MYKL0903 | | 2012 | | Malaysia | N |
| KM055592 | 188A_2007 | | 2007 | | China | N |
| KT359857 | 12MYKL1193 | | 2012 | | Malaysia | N |
| KY862062 | NL63/FRA-EPI/Caen/2012/13 | | 2012 | | France | N |
| KM055596 | 277A_2007 | | 2007 | | China | N |
| KT359831 | 12MYKL0630 | | 2012 | | Malaysia | N |
| KT359865 | 12MYKL1337 | | 2012 | | Malaysia | N |
| KT359858 | 12MYKL1195 | | 2012 | | Malaysia | N |
| KT359852 | 12MYKL1055 | | 2012 | | Malaysia | N |
| KM055600 | 940A_2008 | | 2008 | | China | N |
| KT359848 | 12MYKL0967 | | 2012 | | Malaysia | N |
| KT359835 | 12MYKL0750 | | 2012 | | Malaysia | N |
| KY862056 | NL63/FRA-EPI/Caen/2015/19 | | 2015 | | France | N |
| KT359834 | 12MYKL0736 | | 2012 | | Malaysia | N |
| KT359864 | 12MYKL1336 | | 2012 | | Malaysia | N |
| ***Betacoronavirus***  **Subgenus: *Sarbecovirus*** | | | | | |  |
| **Severe acute respiratory syndrome (SARS)-related coronavirus** | | | | | |  |
| **ID** | **STRAIN** | | **YEAR** | **COUNTRY** | | **ORF** |
| NC_004718.3 | Tor2 | | 2003 | Canada | | Spike, N |
| AP006557 | TWH | | 2003* | Taiwan | | Spike, N |
| AP006558 | TWJ | | 2003* | Taiwan | | Spike, N |
| AP006559 | TWK | | 2003* | Taiwan | | Spike, N |
| AP006560 | TWS | | 2003* | Taiwan | | Spike, N |
| AP006561 | TWY | | 2003* | Taiwan | | Spike, N |
| AY278487 | BJ02 | | 2003 | China | | Spike, N |
| AY278488 | BJ01 | | 2003 | China | | Spike, N |
| AY278489 | GD01 | | 2003 | China | | Spike, N |
| AY278490 | BJ03 | | 2003 | China | | Spike, N |
| AY278491 | HKU-39849 | | 2003 | China | | Spike, N |
| AY278554 | CUHK-W1 | | 2003 | China | | Spike, N |
| AY278741 | Urbani | | 2003 | Atlanta | | Spike, N |
| AY279354 | BJ04 | | 2003 | China | | Spike, N |
| AY282752 | CUHK-Su10 | | 2003 | China | | Spike, N |
| AY291315 | Frankfurt 1 | | 2003 | Germany | | Spike, N |
| AY291451 | TW1 | | 2003 | Taiwan | | Spike, N |
| AY304495 | GZ50 | | 2003 | China | | Spike, N |
| AY310120 | FRA | | 2003 | Germany | | Spike, N |
| AY313906 | GD69 | | 2003 | China | | Spike, N |
| AY321118 | TWC | | 2003* | Taiwan | | Spike, N |
| AY323977 | HSR1 | | 2003 | Italy | | Spike, N |
| AY338174 | TC1 | | 2003* | Taiwan | | Spike, N |
| AY338175 | TC2 | | 2003* | Taiwan | | Spike, N |
| AY345986 | CUHK-AG01 | | 2003 | China | | Spike, N |
| AY345987 | CUHK-AG02 | | 2003 | China | | Spike, N |
| AY345988 | CUHK-AG03 | | 2003 | China | | Spike, N |
| AY348314 | TC3 | | 2003* | Taiwan | | Spike, N |
| AY350750 | PUMC01 | | 2003* | China | | Spike, N |
| AY357075 | PUMC02 | | 2003* | China | | Spike, N |
| AY357076 | PUMC03 | | 2003* | China | | Spike, N |
| AY362698 | TWC2 | | 2003* | Taiwan | | Spike, N |
| AY362699 | TWC3 | | 2003* | Taiwan | | Spike, N |
| AY390556 | GZ02 | | 2003 | China | | Spike, N |
| AY394850 | WHU | | 2004 | China | | Spike, N |
| AY394977 | GZ-A | | 2003 | China | | Spike, N |
| AY394978 | GZ-B | | 2003 | China | | Spike, N |
| AY394979 | GZ-C | | 2003 | China | | Spike, N |
| AY394981 | HGZ8L1-A | | 2003 | China | | Spike, N |
| AY394982 | HGZ8L1-B | | 2003 | China | | Spike, N |
| AY394983 | HSZ2-A | | 2003 | China | | Spike, N |
| AY394984 | HSZ-A | | 2003 | China | | Spike, N |
| AY394985 | HSZ-Bb | | 2003 | China | | Spike, N |
| AY394986 | HSZ-Cb | | 2003 | China | | Spike, N |
| AY394987 | HSZ-Fb | | 2003 | China | | Spike, N |
| AY394988 | JMD | | 2003 | China | | Spike, N |
| AY394989 | HZS2-D | | 2003 | China | | Spike, N |
| AY394990 | HZS2-E | | 2003 | China | | Spike, N |
| AY394991 | HZS2-Fc | | 2003 | China | | Spike, N |
| AY394992 | HZS2-C | | 2003 | China | | Spike, N |
| AY394993 | HGZ8L2 | | 2003 | China | | Spike, N |
| AY394994 | HZS2-Bc | | 2003 | China | | Spike, N |
| AY394995 | HSZ-Cc | | 2003 | China | | Spike, N |
| AY394996 | ZS-B | | 2003 | China | | Spike, N |
| AY394997 | ZS-A | | 2003 | China | | Spike, N |
| AY394998 | LC1 | | 2003 | China | | Spike, N |
| AY394999 | LC2 | | 2003 | China | | Spike, N |
| AY395000 | LC3 | | 2003 | China | | Spike, N |
| AY395001 | LC4 | | 2003 | China | | Spike, N |
| AY395002 | LC5 | | 2003 | China | | Spike, N |
| AY395003 | ZS-C | | 2003 | China | | Spike, N |
| AY395004 | HZS2-Bb | | 2003 | China | | Spike, N |
| AY427439 | AS | | 2003 | Italy | | Spike, N |
| AY461660 | SoD | | 2003* | Russia | | Spike, N |
| AY463059 | ShanghaiQXC1 | | 2003* | China | | Spike, N |
| AY463060 | ShanghaiQXC2 | | 2003* | China | | Spike, N |
| AY485277 | Sino1-11 | | 2003* | China | | Spike, N |
| AY485278 | Sino3-11 | | 2003* | China | | Spike, N |
| AY502923 | TW10 | | 2003 | Taiwan | | Spike, N |
| AY502924 | TW11 | | 2003 | Taiwan | | Spike, N |
| AY502925 | TW2 | | 2003 | Taiwan | | Spike, N |
| AY502926 | TW3 | | 2003 | Taiwan | | Spike, N |
| AY502927 | TW4 | | 2003 | Taiwan | | Spike, N |
| AY502928 | TW5 | | 2003 | Taiwan | | Spike, N |
| AY502929 | TW6 | | 2003 | Taiwan | | Spike, N |
| AY502930 | TW7 | | 2003 | Taiwan | | Spike, N |
| AY502931 | TW8 | | 2003 | Taiwan | | Spike, N |
| AY502932 | TW9 | | 2003 | Taiwan | | Spike, N |
| AY508724 | NS-1 | | 2003* | China | | Spike, N |
| AY559083 | Sin3408 | | 2003* | Singapore | | Spike, N |
| AY559084 | Sin3765V | | 2003* | Singapore | | Spike, N |
| AY559087 | Sin3725V | | 2003* | Singapore | | Spike, N |
| AY595412 | LLJ-2004 | | 2004 | China | | Spike, N |
| AY714217 | CDC#200301157 | | 2003 | USA | | Spike, N |
| AY864806 | BJ202 | | 2004* | China | | Spike, N |
| DQ182595 | ZJ0301 | | 2003 | China | | Spike, N |
| AY864805 | BJ162 | | 2004* | China | | Spike, N |
| EU371559 | ZJ02 | | 2008* | China | | Spike, N |
| DQ640652 | GDH-BJH01 | | 2006* | China | | Spike, N |
| AY297028 | ZJ01 | | 2003* | China | | Spike |
| AH013709 | Sin_WNV | | 2003* | Singapore | | N |
| AY394980 | GZ-D | | 2003 | China | | N |
| AH013708 | Sin0409 | | 2003 | Singapore | | N |
| **Subgenus: *Embecovirus*** | | | | | |  |
| **Human coronavirus OC43** | | | | | | |
| **ID** | **STRAIN** | | **YEAR** | **COUNTRY** | | **ORF** |
| NC_006213 | ATCC VR-759 | | NA | USA | | Spike, N |
| KX538976 | MY-U1057/12 | | 2012 | Malaysia | | Spike, N |
| KY983583 | HCoV_OC43/Seattle/USA/SC2481/2015 | | 2015 | USA | | Spike, N |
| KF530087 | OC43/human/USA/873-6/1987 | | 1987 | USA | | Spike, N |
| KF530081 | OC43/human/USA/991-5/1999 | | 1999 | USA | | Spike, N |
| KF923908 | 5414/2007 | | 2007 | China | | Spike, N |
| KF530095 | OC43/human/USA/912-6/1991 | | 1991 | USA | | Spike, N |
| KF923903 | 12691/2012 | | 2012 | China | | Spike, N |
| AY391777 | OC43 | | NA | NA | | Spike, N |
| KF530072 | OC43/human/USA/9712-13/1997 | | 1997 | USA | | Spike, N |
| MG197711 | BJ-164 | | 2015 | China | | Spike, N |
| KX538966 | MY-U236/12 | | 2012 | Malaysia | | Spike, N |
| KF530082 | OC43/human/USA/912-11/1991 | | 1991 | USA | | Spike, N |
| KF923923 | 892A/2008 | | 2008 | China | | Spike, N |
| KF530093 | OC43/human/USA/832-27/1983 | | 1983 | USA | | Spike, N |
| MK303624 | MDS14 | | NA | NA | | Spike, N |
| MH121121 | HCoV-OC43/USA/ACRI_0213/2016 | | 2016 | USA | | Spike, N |
| MN310476 | HCoV_OC43/Seattle/USA/SC9428/2018 | | 2019 | USA | | Spike, N |
| MG197720 | YC-68 | | 2015 | China | | Spike, N |
| KX538975 | MY-U1024/12 | | 2012 | Malaysia | | Spike, N |
| KF530079 | OC43/human/USA/913-29/1991 | | 1991 | USA | | Spike, N |
| AY585229 | NA | | NA | France | | Spike, N |
| KF530064 | OC43/human/USA/9612-9/1996 | | 1996 | USA | | Spike, N |
| KY967361 | HCoV_OC43/Seattle/USA/SC2345/2015 | | 2015 | USA | | Spike, N |
| KF923922 | 8164/2009 | | 2009 | China | | Spike, N |
| KY369907 | HCoV_OC43/Seattle/USA/SC9741/2016 | | 2016 | USA | | Spike, N |
| KF923907 | 5370/2007 | | 2007 | China | | Spike, N |
| MG977444 | TNP F1778_2 | | 2017 | Cote d'Ivoire | | Spike, N |
| KU131570 | HCoV-OC43/UK/London/2011 | | 2011 | UK | | Spike, N |
| KF530061 | OC43/human/USA/901-43/1990 | | 1990 | USA | | Spike, N |
| MG977452 | TNP 12643 | | 2016 | Cote d'Ivoire | | Spike, N |
| KP198610 | 2058A/10 | | 2010 | China | | Spike, N |
| MG197712 | BJ-165 | | 2015 | China | | Spike, N |
| MK303625 | MDS16 | | NA | NA | | Spike, N |
| MG197716 | WZ-303 | | 2015 | China | | Spike, N |
| KF530065 | OC43/human/USA/901-41/1990 | | 1990 | USA | | Spike, N |
| KF923888 | 2145A/2010 | | 2010 | China | | Spike, N |
| KF923917 | 5566/2007 | | 2007 | China | | Spike, N |
| KY983588 | HCoV_OC43/Seattle/USA/SC3118/2015 | | 2015 | USA | | Spike, N |
| MG977448 | TNP F1833_2 | | 2016 | Cote d'Ivoire | | Spike, N |
| KF530090 | OC43/human/USA/931-85/1993 | | 1993 | USA | | Spike, N |
| MF374983 | HCoV-OC43/USA/TCNP_0070/2016 | | 2016 | USA | | Spike, N |
| KF530096 | OC43/human/USA/911-38/1991 | | 1991 | USA | | Spike, N |
| MG977447 | TNP F1832_2 | | 2016 | Cote d'Ivoire | | Spike, N |
| MN306036 | HCoV_OC43/Seattle/USA/SC0682/2019 | | 2019 | USA | | Spike, N |
| KY014282 | 2007-09 | | 2007 | France | | Spike, N |
| JN129835 | HK04-02 | | 2004 | China | | Spike, N |
| JN129834 | HK04-01 | | 2004 | China | | Spike, N |
| KY967358 | HCoV_OC43/Seattle/USA/SC2770/2015 | | 2015 | USA | | Spike, N |
| KF530060 | OC43/human/USA/851-15/1985 | | 1985 | USA | | Spike, N |
| MG197719 | YC-67 | | 2015 | China | | Spike, N |
| KF530086 | OC43/human/USA/872-5/1987 | | 1987 | USA | | Spike, N |
| KY674917 | N07-1609B | | 2016 | USA | | Spike, N |
| KF923915 | 5517/2007 | | 2007 | China | | Spike, N |
| KX538968 | MY-U464/12 | | 2012 | Malaysia | | Spike, N |
| KF923900 | 3647/2006 | | 2006 | China | | Spike, N |
| KF530076 | OC43/human/USA/911-11/1991 | | 1991 | USA | | Spike, N |
| KF923887 | 1997A/2010 | | 2010 | China | | Spike, N |
| MG977450 | TNP F1835_2 | | 2016 | Cote d'Ivoire | | Spike, N |
| KF923913 | 5485/2007 | | 2007 | China | | Spike, N |
| MF374985 | HCoV-OC43/USA/TCNP_00212/2017 | | 2017 | USA | | Spike, N |
| KY014281 | 2002-04 | | 2002 | France | | Spike, N |
| MK303622 | MDS11 | | NA | NA | | Spike, N |
| MN306041 | HCoV_OC43/Seattle/USA/SC0810/2019 | | 2019 | USA | | Spike, N |
| KF923886 | 1908A/2010 | | 2019 | China | | Spike, N |
| KF923911 | 5479/2007 | | 2007 | China | | Spike, N |
| KF923912 | 5484/2007 | | 2007 | China | | Spike, N |
| KF530098 | OC43/human/USA/965-6/1996 | | 1996 | USA | | Spike, N |
| KF923919 | 5595/2007 | | 2007 | China | | Spike, N |
| KX538977 | MY-U1140/12 | | 2012 | Malaysia | | Spike, N |
| MG197721 | YC-72 | | 2015 | China | | Spike, N |
| KF923892 | 5345/2007 | | 2007 | China | | Spike, N |
| KF530097 | OC43/human/USA/9211-43/1992 | | 1992 | USA | | Spike, N |
| KF923925 | 10574/2010 | | 2010 | China | | Spike, N |
| KF530099 | OC43/human/USA/971-5/1997 | | 1997 | USA | | Spike, N |
| KF530084 | OC43/human/USA/951-18/1995 | | 1995 | USA | | Spike, N |
| KY554975 | N09-382B | | 2016 | USA | | Spike, N |
| KY983585 | HCoV_OC43/Seattle/USA/SC2854/2015 | | 2015 | USA | | Spike, N |
| KF530070 | OC43/human/USA/991-19/1999 | | 1999 | USA | | Spike, N |
| KF923918 | 10108/2010 | | 2010 | China | | Spike, N |
| AY903460 | 19572 Belgium 2004 | | NA | Belgium | | Spike, N |
| KJ958219 | LY342 | | 2011 | China | | Spike, N |
| KF530059 | OC43/human/USA/951-15/1995 | | 1995 | USA | | Spike, N |
| MG977449 | TNP F1834_2 | | 2016 | Cote d'Ivoire | | Spike, N |
| KF530074 | OC43/human/USA/9212-33/1992 | | 1992 | USA | | Spike, N |
| KF923902 | 12689/2012 | | 2012 | China | | Spike, N |
| KF923893 | 2151A/2010 | | 2010 | China | | Spike, N |
| KY684759 | HCoV_OC43/Seattle/USA/SC2269/2016 | | 2016 | USA | | Spike, N |
| MG197717 | WZ-522 | | 2015 | China | | Spike, N |
| KF530071 | OC43/human/USA/925-1/1992 | | 1992 | USA | | Spike, N |
| MK303619 | MDS6 | | NA | NA | | Spike, N |
| KF530073 | OC43/human/USA/8912-37/1989 | | 1989 | USA | | Spike, N |
| KX538965 | MY-U208/12 | | 2012 | Malaysia | | Spike, N |
| MK327281 | MDS15 | | NA | NA | | Spike, N |
| KF530083 | OC43/human/USA/873-19/1987 | | 1987 | USA | | Spike, N |
| MN306042 | HCoV_OC43/Seattle/USA/SC0839/2019 | | 2019 | USA | | Spike, N |
| KF530063 | OC43/human/USA/9612-48/1996 | | 1996 | USA | | Spike, N |
| MK303621 | MDS4 | | NA | NA | | Spike, N |
| KF923896 | 3074A/2012 | | 2012 | China | | Spike, N |
| KX538973 | MY-U868/12 | | 2012 | Malaysia | | Spike, N |
| KF923898 | 3184A/2012 | | 2012 | China | | Spike, N |
| MG977446 | TNP F1791_2 | | 2016 | Cote d'Ivoire | | Spike, N |
| MG977451 | TNP 12636 | | 2016 | Cote d'Ivoire | | Spike, N |
| MG197714 | CC-23 | | 2015 | China | | Spike, N |
| KF530080 | OC43/human/USA/9712-31/1997 | | 1997 | USA | | Spike, N |
| KF530092 | OC43/human/USA/008-5/2000 | | 2000 | USA | | Spike, N |
| KY967356 | HCoV_OC43/Seattle/USA/SC2924/2015 | | 2015 | USA | | Spike, N |
| KY967360 | HCoV_OC43/Seattle/USA/SC2476/2015 | | 2015 | USA | | Spike, N |
| MG197715 | GZYF-26 | | 2015 | China | | Spike, N |
| MN310478 | HCoV_OC43/Seattle/USA/SC0776/2019 | | 2019 | USA | | Spike, N |
| KX538969 | MY-U523/12 | | 2012 | Malaysia | | Spike, N |
| KF530078 | OC43/human/USA/9612-29/1996 | | 1996 | USA | | Spike, N |
| KY369905 | HCoV_OC43/Seattle/USA/SC831/2016 | | 2016 | USA | | Spike, N |
| MG197709 | BJ-112 | | 2015 | China | | Spike, N |
| KY554973 | N07-1689B_116X | | 2016 | USA | | Spike, N |
| KF923894 | 5352/2007 | | 2007 | China | | Spike, N |
| MG197718 | YC-55 | | 2015 | China | | Spike, N |
| KX538979 | MY-U1975/13 | | 2013 | Malaysia | | Spike, N |
| MG197713 | BJ-221 | | 2015 | China | | Spike, N |
| KX538978 | MY-U1758/13 | | 2013 | Malaysia | | Spike, N |
| KF923924 | 10290/2010 | | 2010 | China | | Spike, N |
| KF530077 | OC43/human/USA/873-16/1987 | | 1987 | USA | | Spike, N |
| KF923895 | 10285/2010 | | 2010 | China | | Spike, N |
| KX344031 | LRTI_238 | | 2011 | Mexico | | Spike, N |
| KF923890 | 39A/2007 | | 2007 | China | | Spike, N |
| KY674918 | N07-1647B | | 2016 | USA | | Spike, N |
| KF923916 | 5519/2007 | | 2007 | China | | Spike, N |
| KF530062 | OC43/human/USA/952-23/1995 | | 1995 | USA | | Spike, N |
| KF530085 | OC43/human/USA/871-25/1987 | | 1987 | USA | | Spike, N |
| KY967359 | HCoV_OC43/Seattle/USA/SC2730/2015 | | 2015 | USA | | Spike, N |
| KF530089 | OC43/human/USA/911-66/1991 | | 1991 | USA | | Spike, N |
| MF374984 | HCoV-OC43/USA/TCNP_00204/2017 | | 2017 | USA | | Spike, N |
| KF923905 | 229/2005 | | 2005 | China | | Spike, N |
| KF923909 | 5442/2007 | | 2007 | China | | Spike, N |
| KX538971 | MY-U732/12 | | 2012 | Malaysia | | Spike, N |
| KX538972 | MY-U774/12 | | 2012 | Malaysia | | Spike, N |
| KX538967 | MY-U413/12 | | 2012 | Malaysia | | Spike, N |
| KF923910 | 5445/2007 | | 2007 | China | | Spike, N |
| KF530069 | OC43/human/USA/982-4/1998 | | 1998 | USA | | Spike, N |
| MG197710 | BJ-124 | | 2015 | China | | Spike, N |
| KF923920 | 5617/2007 | | 2007 | China | | Spike, N |
| KF530088 | OC43/human/USA/901-54/1990 | | 1990 | USA | | Spike, N |
| KY554972 | N07-1541B_433X | | 2016 | USA | | Spike, N |
| KY674920 | N09-595B | | 2016 | USA | | Spike, N |
| KX538970 | MY-U710/12 | | 2012 | Malaysia | | Spike, N |
| KF923891 | 5240/2007 | | 2007 | China | | Spike, N |
| KF530094 | OC43/human/USA/912-36/1991 | | 1991 | USA | | Spike, N |
| KF923899 | 3582/2006 | | 2006 | China | | Spike, N |
| KF923906 | 3194A/2012 | | 2012 | China | | Spike, N |
| KF530067 | OC43/human/USA/912-10/1991 | | 1991 | USA | | Spike, N |
| MN026164 | OC43_KLF_01_2018 | | 2018 | Kenya | | Spike, N |
| MG197722 | YC-207 | | 2015 | China | | Spike, N |
| KX538964 | MY-U002/12 | | 2015 | Malaysia | | Spike, N |
| KF923921 | 69A/2007 | | 2007 | China | | Spike, N |
| KF923897 | 3269A/2012 | | 2012 | China | | Spike, N |
| KY554974 | N08-33B_360X | | 2016 | USA | | Spike, N |
| KF530091 | OC43/human/USA/911-58/1991 | | 1991 | USA | | Spike, N |
| MG197723 | HZ-459 | | 2016 | China | | Spike, N |
| KF530066 | OC43/human/USA/901-33/1990 | | 1990 | USA | | Spike, N |
| KX538974 | MY-U945/12 | | 2012 | Malaysia | | Spike, N |
| MG977445 | TNP F1790_2 | | 2016 | Cote d'Ivoire | | Spike, N |
| KJ958218 | LY341 | | 2011 | China | | Spike, N |
| KF923914 | 5508/2007 | | 2007 | China | | Spike, N |
| AY903459 | 87309 | | NA | Belgium | | Spike, N |
| KF923901 | 5472/2007 | | 2007 | China | | Spike, N |
| MN306043 | HCoV_OC43/Seattle/USA/SC0841/2019 | | 2019 | USA | | Spike, N |
| KF530068 | OC43/human/USA/007-11/2000 | | 2000 | USA | | Spike, N |
| MK303620 | MDS2 | | NA | NA | | Spike, N |
| MN306053 | HCoV_OC43/Seattle/USA/SC9430/2018 | | 2019 | USA | | Spike, N |
| KF923889 | 1926/2006 | | 2006 | China | | Spike, N |
| KF923904 | 12694/2012 | | 2012 | China | | Spike, N |
| KP198611 | 1783A/10 | | 2010 | China | | Spike, N |
| KF530075 | OC43/human/USA/953-23/1995 | | 2010 | USA | | Spike, N |
| KY369906 | HCoV_OC43/Seattle/USA/SC622/2016 | | 2016 | USA | | Spike, N |
| MF314143 | HCoV-OC43/USA/ACRI_0052/2016 | | 2016 | USA | | Spike, N |
| KF572805 | 1908A_10 | | 2010 | China | | Spike |
| KF572825 | 10574_10 | | 2010 | China | | Spike |
| KF572852 | 5472_07 | | 2007 | China | | Spike |
| KF572850 | 5442_07 | | 2007 | China | | Spike |
| KF572824 | 1034A_08 | | 2008 | China | | Spike |
| L14643 | OC43 | | NA | NA | | Spike |
| LC315647 | Tokyo/SGH-61/2014 | | 2014 | Japan | | Spike |
| KU745536 | 14012/2014 | | 2014 | China | | Spike |
| KF572831 | 12689_12 | | 2012 | China | | Spike |
| LC315646 | Tokyo/SGH-36/2014 | | 2014 | Japan | | Spike |
| KF572816 | 229_05 | | 2005 | China | | Spike |
| AY903458 | 36638 Belgium 2004 | | NA | Belgium | | Spike |
| KF572864 | 69A_07 | | 2007 | China | | Spike |
| KF572844 | 5240_07 | | 2007 | China | | Spike |
| KF572833 | 12694_12 | | 2012 | China | | Spike |
| KF572862 | 5625_07 | | 2007 | China | | Spike |
| KF572853 | 5479_07 | | 2007 | China | | Spike |
| AY903455 | 34364 Belgium 2004 | | NA | Belgium | | Spike |
| KF572838 | 2134A_10 | | 2010 | China | | Spike |
| KF572872 | 978A_08 | | 2008 | China | | Spike |
| KU745537 | 3791A/2013 | | 2013 | China | | Spike |
| KF572868 | 892A_08 | | 2008 | China | | Spike |
| KF572814 | 3194A_12 | | 2012 | China | | Spike |
| KU745543 | 4449A/2015 | | 2015 | China | | Spike |
| KU745544 | 4450A/2015 | | 2015 | China | | Spike |
| KU745545 | 4452A/2015 | | 2015 | China | | Spike |
| KF572830 | 1216A_08 | | 2008 | China | | Spike |
| KF572865 | 8099_09 | | 2009 | China | | Spike |
| KF572857 | 5517_07 | | 2007 | China | | Spike |
| KF572861 | 5617_07 | | 2007 | China | | Spike |
| KF572811 | 2941A_11 | | 2011 | China | | Spike |
| KF572846 | 5345_07 | | 2007 | China | | Spike |
| KF572826 | 1081A_08 | | 2008 | China | | Spike |
| KF572809 | 2058A_10 | | 2010 | China | | Spike |
| KF572810 | 2145A_10 | | 2010 | China | | Spike |
| KF572822 | 10285_10 | | 2010 | China | | Spike |
| KU745538 | 4068A/2014 | | 2014 | China | | Spike |
| KU745534 | 13969/2014 | | 2014 | China | | Spike |
| KF572828 | 1157A_08 | | 2008 | China | | Spike |
| KF963236 | HCoV-OC43/FRA_EPI/Caen/2005/07 | | 2005 | France | | Spike |
| KF572843 | 4954_07 | | 2007 | China | | Spike |
| KF572829 | 11930_11 | | 2011 | China | | Spike |
| KF572839 | 2151A_10 | | 2010 | China | | Spike |
| KF572812 | 3074A_12 | | 2012 | China | | Spike |
| KF572845 | 5331_07 | | 2007 | China | | Spike |
| KF963235 | HCoV-OC43/FRA_EPI/Caen/2004/06 | | 2004 | France | | Spike |
| KF963243 | HCoV-OC43/FRA_EPI/Caen/2012/14 | | 2012 | France | | Spike |
| KF572854 | 5484_07 | | 2007 | China | | Spike |
| KF572866 | 8164_09 | | 2009 | China | | Spike |
| KU745535 | 14007/2014 | | 2014 | China | | Spike |
| KF572842 | 4795_07 | | 2007 | China | | Spike |
| KF963240 | HCoV-OC43/FRA_EPI/Caen/2009/11 | | 2009 | France | | Spike |
| KU745548 | SZ14014d3/2014 | | 2014 | China | | Spike |
| AY903457 | 37767 Belgium 2003 | | NA | Belgium | | Spike |
| KF572815 | 1489A_09 | | 2009 | China | | Spike |
| KF572847 | 5352_07 | | 2007 | China | | Spike |
| KF572806 | 1919A_10 | | 2010 | China | | Spike |
| KF963244 | HCoV-OC43/FRA_EPI/Caen/2013/15 | | 2013 | France | | Spike |
| KF963229 | HCoV-OC43/FRA_EPI/Caen/1967/VR759 | | 1967 | France | | Spike |
| KU745533 | 13963/2014 | | 2014 | China | | Spike |
| KF572840 | 3098A_12 | | 2012 | China | | Spike |
| KF572817 | 3582_06 | | 2006 | China | | Spike |
| KF572813 | 3184A_12 | | 2012 | China | | Spike |
| KF572856 | 5508_07 | | 2007 | China | | Spike |
| KF963238 | HCoV-OC43/FRA_EPI/Caen/2007/09 | | 2007 | France | | Spike |
| KF572855 | 5485_07 | | 2007 | China | | Spike |
| KU745539 | 4086A/2014 | | 2014 | China | | Spike |
| KF572818 | 3647_06 | | 2006 | China | | Spike |
| KF572819 | 039A_07 | | 2007 | China | | Spike |
| KF572851 | 5445_07 | | 2007 | China | | Spike |
| KF572808 | 1997A_10 | | 2010 | China | | Spike |
| KF963230 | HCoV-OC43/FRA_EPI/Caen/2001/01 | | 2001 | France | | Spike |
| KF963231 | HCoV-OC43/FRA_EPI/Caen/2001/02 | | 2001 | France | | Spike |
| KU745540 | 4400A/2015 | | 2015 | China | | Spike |
| KF572821 | 10108_10 | | 2010 | China | | Spike |
| KF963232 | HCoV-OC43/FRA_EPI/Caen/2002/03 | | 2002 | France | | Spike |
| KF572823 | 10290_10 | | 2010 | China | | Spike |
| KF572804 | 1783A_10 | | 2010 | China | | Spike |
| KU745547 | HB14018d7/2014 | | 2014 | China | | Spike |
| KF572869 | 8942_09 | | 2009 | China | | Spike |
| KF963242 | HCoV-OC43/FRA_EPI/Caen/2011/13 | | 2011 | France | | Spike |
| KF572835 | 1382A_09 | | 2009 | China | | Spike |
| KF572832 | 12691_12 | | 2012 | China | | Spike |
| AY903456 | 84020 Belgium 2003 | | NA | Belgium | | Spike |
| KF963233 | HCoV-OC43/FRA_EPI/Caen/2002/04 | | 2002 | France | | Spike |
| KF572859 | 5566_07 | | 2007 | China | | Spike |
| KU745541 | 4436A/2015 | | 2015 | China | | Spike |
| KF572827 | 1135A_08 | | 2008 | China | | Spike |
| KU745546 | 4467A/2015 | | 2015 | China | | Spike |
| KF572849 | 5414_07 | | 2007 | China | | Spike |
| Z21849 | HCV-OC43 | | NA | NA | | Spike |
| KF572807 | 1926_06 | | 2006 | China | | Spike |
| KF572848 | 5370_07 | | 2007 | China | | Spike |
| KF963241 | HCoV-OC43/FRA_EPI/Caen/2010/12 | | 2010 | France | | Spike |
| KF572836 | 1591A_09 | | 2009 | China | | Spike |
| KF572867 | 8375_09 | | 2009 | China | | Spike |
| KF572863 | 5656_07 | | 2007 | China | | Spike |
| KF572841 | 3269A_12 | | 2012 | China | | Spike |
| KF572870 | 9001_09 | | 2009 | China | | Spike |
| KF572860 | 5595_07 | | 2007 | China | | Spike |
| KF572837 | 1593A_09 | | 2009 | China | | Spike |
| KF572820 | 079A_07 | | 2007 | China | | Spike |
| AY903454 | 89996 Belgium 2003 | | NA | Belgium | | Spike |
| KU745542 | 4446A/2015 | | 2015 | China | | Spike |
| KF572858 | 5519_07 | | 2007 | China | | Spike |
| KF572834 | 1357A_09 | | 2009 | China | | Spike |
| KF963239 | HCoV-OC43/FRA_EPI/Caen/2008/10 | | 2008 | France | | Spike |
| KF572871 | 9138_09 | | 2009 | China | | Spike |
| LC315648 | Tokyo/SGH-06/2015 | | 2015 | Japan | | Spike |
| KF963234 | HCoV-OC43/FRA_EPI/Caen/2003/05 | | 2003 | France | | Spike |
| KF963237 | HCoV-OC43/FRA_EPI/Caen/2006/08 | | 2006 | France | | Spike |
| LC315649 | Tokyo/SGH-65/2016 | | 2016 | Japan | | Spike |
| KR055616 | 12MYKL1140 | | 2012 | Malaysia | | N |
| KF572739 | 1783A_10 | | 2010 | China | | N |
| KU745579 | HB14018d7/2014 | | 2014 | China | | N |
| KF963200 | HCoV-OC43/FRA_EPI/Caen/2002/03 | | 2002 | France | | N |
| KR055605 | 12MYKL0523 | | 2012 | Malaysia | | N |
| KF963207 | HCoV-OC43/FRA_EPI/Caen/2008/10 | | 2008 | France | | N |
| KU745573 | 4436A/2015 | | 2015 | China | | N |
| KF572799 | 9001_09 | | 2009 | China | | N |
| KF572800 | 9138_09 | | 2009 | China | | N |
| KF572789 | 5656_07 | | 2007 | China | | N |
| KF572750 | 2145A_10 | | 2010 | China | | N |
| KF963211 | HCoV-OC43/FRA_EPI/Caen/2012/14 | | 2012 | France | | N |
| KF572786 | 5617_07 | | 2007 | China | | N |
| KF572708 | 079A_07 | | 2007 | China | | N |
| KF572722 | 11930_11 | | 2011 | China | | N |
| KF572710 | 10241_10 | | 2010 | China | | N |
| KF572736 | 1591A_09 | | 2009 | China | | N |
| KF572798 | 8942_09 | | 2009 | China | | N |
| KR055620 | 13MYKL1758 | | 2013 | Malaysia | | N |
| KF572740 | 1908A_10 | | 2010 | China | | N |
| KF572721 | 11914_11 | | 2011 | China | | N |
| KF572751 | 2151A_10 | | 2010 | China | | N |
| KF572735 | 1537A_09 | | 2009 | China | | N |
| KF572709 | 10108_10 | | 2010 | China | | N |
| KF572743 | 1996A_10 | | 2010 | China | | N |
| KF572758 | 3582_06 | | 2006 | China | | N |
| KF572797 | 892A_08 | | 2008 | China | | N |
| KF572780 | 5519_07 | | 2007 | China | | N |
| KF572795 | 8164_09 | | 2009 | China | | N |
| KF572753 | 229_05 | | 2005 | China | | N |
| KU745576 | 4450A/2015 | | 2015 | China | | N |
| KF572760 | 3647_06 | | 2006 | China | | N |
| KF572746 | 3074A_12 | | 2012 | China | | N |
| KR055601 | 12MYKL0208 | | 2012 | Malaysia | | N |
| KF572725 | 12689_12 | | 2012 | China | | N |
| KR055619 | 12MYKL1612 | | 2012 | Malaysia | | N |
| KR055599 | 12MYKL0002 | | 2012 | Malaysia | | N |
| KF572773 | 5462_07 | | 2007 | China | | N |
| KF572796 | 8375_09 | | 2009 | China | | N |
| KF572776 | 5484_07 | | 2007 | China | | N |
| KF963209 | HCoV-OC43/FRA_EPI/Caen/2010/12 | | 2010 | France | | N |
| KF572748 | 3194A_10 | | 2010 | China | | N |
| KR055617 | 12MYKL1381 | | 2012 | Malaysia | | N |
| KU745567 | 14007/2014 | | 2014 | China | | N |
| KF572719 | 1155A_08 | | 2008 | China | | N |
| KF572757 | 3269A_10 | | 2012 | China | | N |
| KF572767 | 5352_07 | | 2007 | China | | N |
| KF572787 | 5625_07 | | 2007 | China | | N |
| KF572791 | 648_05 | | 2005 | China | | N |
| KF963210 | HCoV-OC43/FRA_EPI/Caen/2011/13 | | 2011 | France | | N |
| KF572794 | 8099_09 | | 2009 | China | | N |
| KF572754 | 3089A_10 | | 2010 | China | | N |
| KF572803 | O75A_07 | | 2007 | China | | N |
| KF572730 | 12747_12 | | 2012 | China | | N |
| KF572793 | 7712_08 | | 2008 | China | | N |
| KF572774 | 5472_07 | | 2007 | China | | N |
| KF572765 | 5331_07 | | 2007 | China | | N |
| KF572733 | 1382A_09 | | 2009 | China | | N |
| KF572781 | 5566_07 | | 2007 | China | | N |
| KF572802 | 9817_10 | | 2010 | China | | N |
| KF963203 | HCoV-OC43/FRA_EPI/Caen/2004/06 | | 2004 | France | | N |
| KF572711 | 10285_10 | | 2010 | China | | N |
| KF572741 | 1919A_10 | | 2010 | China | | N |
| KU745565 | 13963/2014 | | 2014 | China | | N |
| KF572714 | 1034A_08 | | 2008 | China | | N |
| KF572716 | 10574_10 | | 2010 | China | | N |
| KF572727 | 12694_12 | | 2012 | China | | N |
| KU745570 | 4068A/2014 | | 2014 | China | | N |
| KR055612 | 12MYKL0945 | | 2012 | Malaysia | | N |
| KF572726 | 12691_12 | | 2012 | China | | N |
| KF572717 | 1081A_08 | | 2008 | China | | N |
| KF572778 | 5508_07 | | 2007 | China | | N |
| KR055610 | 12MYKL0781 | | 2012 | Malaysia | | N |
| KF963205 | HCoV-OC43/FRA_EPI/Caen/2006/08 | | 2006 | France | | N |
| KF963201 | HCoV-OC43/FRA_EPI/Caen/2002/04 | | 2002 | France | | N |
| KF572762 | 4795_07 | | 2007 | China | | N |
| KF572720 | 1157A_08 | | 2008 | China | | N |
| KF572763 | 4954_07 | | 2007 | China | | N |
| KF572715 | 10352_10 | | 2010 | China | | N |
| KF572756 | 3184A_10 | | 2010 | China | | N |
| KF572770 | 5442_07 | | 2007 | China | | N |
| KF572752 | 2171A_10 | | 2010 | China | | N |
| KR055602 | 12MYKL0236 | | 2012 | Malaysia | | N |
| KU745578 | 4467A/2015 | | 2015 | China | | N |
| KU745569 | 3791A/2013 | | 2013 | China | | N |
| KF572777 | 5485_07 | | 2007 | China | | N |
| KF963197 | HCoV-OC43/FRA_EPI/Caen/1967/VR759 | | 1967 | France | | N |
| KU745577 | 4452A/2015 | | 2015 | China | | N |
| KF963212 | HCoV-OC43/FRA_EPI/Caen/2013/15 | | 2015 | France | | N |
| KR055613 | 12MYKL1024 | | 2012 | Malaysia | | N |
| KF572745 | 2058A_10 | | 2020 | China | | N |
| KU745572 | 4400A/2015 | | 2015 | China | | N |
| KF572731 | 12760_12 | | 2012 | China | | N |
| KF572744 | 1997A_10 | | 2010 | China | | N |
| KF572785 | 5595_07 | | 2007 | China | | N |
| KF572784 | 5570_07 | | 2007 | China | | N |
| KR055614 | 12MYKL1130 | | 2012 | Malaysia | | N |
| KF963204 | HCoV-OC43/FRA_EPI/Caen/2005/07 | | 2005 | France | | N |
| KF572772 | 5448_07 | | 2007 | China | | N |
| KR055603 | 12MYKL0413 | | 2012 | Malaysia | | N |
| KF572771 | 5445_07 | | 2007 | China | | N |
| KF572768 | 5370_07 | | 2007 | China | | N |
| KF572729 | 12741_12 | | 2012 | China | | N |
| KF572779 | 5517_07 | | 2007 | China | | N |
| KF572792 | 69A_07 | | 2007 | China | | N |
| KF572712 | 10290_10 | | 2010 | China | | N |
| KU745571 | 4086A/2014 | | 2014 | China | | N |
| KF572801 | 978A_08 | | 2008 | China | | N |
| KF572755 | 3098A_10 | | 2010 | China | | N |
| KF572764 | 5240_07 | | 2007 | China | | N |
| KF963198 | HCoV-OC43/FRA_EPI/Caen/2001/01 | | 2001 | France | | N |
| KF963208 | HCoV-OC43/FRA_EPI/Caen/2009/11 | | 2009 | France | | N |
| KF572738 | 1702_06 | | 2006 | China | | N |
| KR055621 | 13MYKL1975 | | 2013 | Malaysia | | N |
| KU745580 | SZ14014d3/2014 | | 2014 | China | | N |
| KF572734 | 1489A_09 | | 2009 | China | | N |
| KF963206 | HCoV-OC43/FRA_EPI/Caen/2007/09 | | 2007 | France | | N |
| KF572759 | 3630_06 | | 2006 | China | | N |
| KR055608 | 12MYKL0732 | | 2012 | Malaysia | | N |
| KF572718 | 1135A_08 | | 2008 | China | | N |
| KF572724 | 12651_12 | | 2012 | China | | N |
| KU745575 | 4449A/2015 | | 2015 | China | | N |
| KR055611 | 12MYKL0868 | | 2012 | Malaysia | | N |
| KU745566 | 13969/2014 | | 2014 | China | | N |
| KF572728 | 12700_12 | | 2012 | China | | N |
| KF572761 | 39A_07 | | 2007 | China | | N |
| KF572723 | 1216A_08 | | 2008 | China | | N |
| KR055606 | 12MYKL0710 | | 2012 | Malaysia | | N |
| KF572783 | 5569_07 | | 2007 | China | | N |
| KU745574 | 4446A/2015 | | 2015 | China | | N |
| KF572790 | 607_05 | | 2005 | China | | N |
| KU745568 | 14012/2014 | | 2014 | China | | N |
| KF572749 | 2134A_10 | | 2010 | China | | N |
| KF963199 | HCoV-OC43/FRA_EPI/Caen/2001/02 | | 2001 | France | | N |
| KF572766 | 5345_07 | | 2007 | China | | N |
| KF572747 | 2941A_11 | | 2011 | China | | N |
| KR055600 | 12MYKL0043 | | 2012 | Malaysia | | N |
| KF572769 | 5414_07 | | 2007 | China | | N |
| KF572713 | 1029A_08 | | 2008 | China | | N |
| KR055615 | 12MYKL1057 | | 2012 | Malaysia | | N |
| KF572742 | 1926_06 | | 2006 | China | | N |
| KF963202 | HCoV-OC43/FRA_EPI/Caen/2003/05 | | 2003 | France | | N |
| KF572775 | 5479_07 | | 2007 | China | | N |
| KR055618 | 12MYKL1484 | | 2012 | Malaysia | | N |
| KF572788 | 5652_07 | | 2007 | China | | N |
| KF572782 | 5567_07 | | 2007 | China | | N |
| KF572732 | 1357A_09 | | 2009 | China | | N |
| KF572737 | 1593A_09 | | 2009 | China | | N |
| KR055607 | 12MYKL0774 | | 2012 | Malaysia | | N |
| KR055609 | 12MYKL0760 | | 2012 | Malaysia | | N |
| KR055604 | 12MYKL0464 | | 2012 | Malaysia | | N |
| **Human coronavirus HKU1** | | | | | | |
| **ID** | | **STRAIN** | **YEAR** | **COUNTRY** | | **ORFs** |
| NC_006577 | | HKU1 | NA | NA | | Spike, N |
| KF430201 | | HKU1/human/USA/HKU1-18/2010 | 2010 | USA | | Spike, N |
| DQ415901 | | N24 | NA | China | | Spike, N |
| KF686341 | | HKU1/human/USA/HKU1-10/2010 | 2010 | USA | | Spike, N |
| DQ415908 | | N11 | NA | China | | Spike, N |
| KF686343 | | HKU1/human/USA/HKU1-13/2010 | 2010 | USA | | Spike, N |
| KF686344 | | HKU1/human/USA/HKU1-15/2009 | 2009 | USA | | Spike, N |
| DQ415903 | | N3 | NA | China | | Spike, N |
| DQ415905 | | N7 | NA | China | | Spike, N |
| KY674942 | | N09-1627B | 2016 | USA | | Spike, N |
| KF686339 | | HKU1/human/USA/HKU1-3/2009 | 2009 | USA | | Spike, N |
| KY674921 | | N08-87 | 2016 | USA | | Spike, N |
| MH940245 | | SI17244 | 2017 | Thailand | | Spike, N |
| DQ415902 | | N25 | NA | China | | Spike, N |
| DQ415909 | | N13 | NA | China | | Spike, N |
| DQ415899 | | N22 | NA | China | | Spike, N |
| DQ415910 | | N14 | NA | China | | Spike, N |
| DQ415913 | | N17 | NA | China | | Spike, N |
| KF686340 | | HKU1/human/USA/HKU1-5/2009 | 2009 | USA | | Spike, N |
| KF686342 | | HKU1/human/USA/HKU1-11/2009 | 2009 | USA | | Spike, N |
| DQ339101 | | N5P8 | NA | NA | | Spike, N |
| KF686346 | | HKU1/human/USA/HKU1-12/2010 | 2010 | USA | | Spike, N |
| DQ415907 | | N10 | NA | China | | Spike, N |
| KT779556 | | BJ01-p9 | 2009 | China | | Spike, N |
| KY983584 | | SC2628 | 2015 | USA | | Spike, N |
| KF686338 | | HKU1/human/USA/HKU1-1/2005 | 2005 | USA | | Spike, N |
| DQ415898 | | N21 | NA | China | | Spike, N |
| DQ415911 | | N15 | NA | China | | Spike, N |
| AY884001 | | NA | NA | NA | | Spike, N |
| KF430199 | | HKU1/human/USA/HKU1-14/2009 | 2009 | USA | | Spike, N |
| KF686345 | | HKU1/human/USA/HKU1-20/2010 | 2010 | USA | | Spike, N |
| KF430200 | | HKU1/human/USA/HKU1-16/2010 | 2010 | USA | | Spike, N |
| DQ415896 | | N19 | NA | China | | Spike, N |
| KY674943 | | N09-1605B | 2016 | USA | | Spike, N |
| DQ415897 | | N20 | NA | China | | Spike, N |
| MK167038 | | SC2521 | 2017 | USA | | Spike, N |
| DQ415906 | | N9 | NA | China | | Spike, N |
| KY674941 | | N09-1663B | 2016 | USA | | Spike, N |
| KT779555 | | BJ01-p3 | 2009 | China | | Spike, N |
| HM034837 | | Caen1 | 2005 | France | | Spike, N |
| DQ415900 | | N23 | NA | China | | Spike, N |
| DQ415914 | | N18 | NA | China | | Spike, N |
| DQ415904 | | N6 | NA | China | | Spike, N |
| KF430202 | | HKU1/human/USA/HKU1-7/2010 | 2010 | USA | | Spike, N |
| DQ415912 | | N16 | NA | China | | Spike, N |
| DQ437619 | | N24 | NA | Hong Kong | | Spike |
| DQ437615 | | N20 | NA | Hong Kong | | Spike |
| KF430203 | | HKU1/HTBEC_lab/BRA/HKU1-23/2006 | 2006 | Brazil | | Spike |
| DQ437609 | | N14 | NA | Hong Kong | | Spike |
| DQ437611 | | N16 | NA | Hong Kong | | Spike |
| DQ437613 | | N18 | NA | Hong Kong | | Spike |
| DQ437614 | | N19 | NA | Hong Kong | | Spike |
| DQ437618 | | N23 | NA | Hong Kong | | Spike |
| KF430197 | | HKU1/HTBEC_lab/BRA/HKU1-22/2007 | 2007 | Brazil | | Spike |
| DQ437607 | | N11 | NA | Hong Kong | | Spike |
| KF430198 | | HKU1/HTBEC_lab/BRA/HKU1-21/2006 | 2006 | Brazil | | Spike |
| DQ437612 | | N17 | NA | Hong Kong | | Spike |
| DQ437616 | | N21 | NA | Hong Kong | | Spike |
| DQ437610 | | N15 | NA | Hong Kong | | Spike |
| DQ437617 | | N22 | NA | Hong Kong | | Spike |
| DQ437608 | | N13 | NA | Hong Kong | | Spike |
| LC315650 | | Tokyo/SGH-15/2014 | 2014 | Japan | | Spike |
| LC315651 | | Tokyo/SGH-18/2016 | 2016 | Japan | | Spike |
| KR055537 | | 12MYKL0529 | 2012 | Malaysia | | N |
| KR055532 | | 12MYKL0163 | 2012 | Malaysia | | N |
| KR055540 | | 12MYKL0759 | 2012 | Malaysia | | N |
| DQ437621 | | N13 | NA | Hong Kong | | N |
| DQ437632 | | N24 | NA | Hong Kong | | N |
| KR055547 | | 12MYKL1132 | 2012 | Malaysia | | N |
| KR055541 | | 12MYKL0777 | 2012 | Malaysia | | N |
| KR055534 | | 12MYKL0323 | 2012 | Malaysia | | N |
| KR055551 | | 13MYKL1781 | 2013 | Malaysia | | N |
| DQ437623 | | N15 | NA | Hong Kong | | N |
| KR055533 | | 12MYKL0181 | 2012 | Malaysia | | N |
| KR055545 | | 12MYKL1061 | 2012 | Malaysia | | N |
| KF850450 | | HKU1 1102 | 2005 | USA | | N |
| KR055546 | | 12MYKL1075 | 2012 | Malaysia | | N |
| KR055544 | | 12MYKL1058 | 2012 | Malaysia | | N |
| KR055550 | | 12MYKL1217 | 2012 | Malaysia | | N |
| KR055539 | | 12MYKL0737 | 2012 | Malaysia | | N |
| KR055552 | | 13MYKL1898 | 2013 | Malaysia | | N |
| DQ437629 | | N21 | NA | Hong Kong | | N |
| KR055548 | | 12MYKL1153 | 2012 | Malaysia | | N |
| DQ437631 | | N23 | NA | Hong Kong | | N |
| KR055535 | | 12MYKL0407 | 2012 | Malaysia | | N |
| KR055538 | | 12MYKL0624 | 2012 | Malaysia | | N |
| DQ437628 | | N20 | NA | Hong Kong | | N |
| DQ437625 | | N17 | NA | Hong Kong | | N |
| DQ778921 | | Caen | NA | France | | N |
| KR055549 | | 12MYKL1214 | 2012 | Malaysia | | N |
| KF430196 | | HKU1/human/USA/HKU1-4/2005 | 2005 | USA | | N |
| DQ437624 | | N16 | NA | Hong Kong | | N |
| DQ437627 | | N19 | NA | Hong Kong | | N |
| DQ437620 | | N11 | NA | Hong Kong | | N |
| KR055542 | | 12MYKL0790 | 2012 | Malaysia | | N |
| KR055543 | | 12MYKL0841 | 2012 | Malaysia | | N |
| DQ437622 | | N14 | NA | Hong Kong | | N |
| KR055536 | | 12MYKL0447 | 2012 | Malaysia | | N |
| DQ437630 | | N22 | NA | Hong Kong | | N |
| KR055553 | | 13MYKL1997 | 2013 | Malaysia | | N |
| DQ437626 | | N18 | NA | Hong Kong | | N |
| *Deposition date | | | | | | |
