## Supplementary text 1 for "Antigenic variation of SARS-CoV-2 in response to immune pressure"

### For each protein we evaluated the difference D between average H values at epitope and non-epitope positions. 
### Most positions of analyzed viruses are invariable along the alignments, so the distribution of H is zero-inflated. 
### We thus calculated statistical significance by permutations.  For each protein, the predicted epitope intervals were 
### collapsed to a single position while non-epitope intervals were left unchanged. 
### After randomly shuffling this collapsed sequence it was expanded back to full length and the difference between 
### shuffled epitope and non-epitope H values was calculated.  
### This procedure was repeated 1000 times and the proportion of repetitions showing a difference more extreme than D 
### was reported as p-value. 

permuteEpitopes<-function(epitopeAnnotation,entropy,N=1000){

  collapseEpitope<-function(epi){
    posprotein<-1;
    poslista<-0;
    L<-list();
    prev<-0
    while(posprotein<=length(epi)){
      if(is.na(epi[posprotein])){
        poslista<-poslista+1;
        L[[poslista]]<-NA;
      }else if(epi[posprotein]==0 & !is.na(epi[posprotein])){
        poslista<-poslista+1;
        L[[poslista]]<-0;
      }else if (epi[posprotein]==1 & !is.na(epi[posprotein])){
        if(prev==0 | is.na(prev) ){
          poslista<-poslista+1
        }
        if(poslista>length(L)){
          L[[poslista]]<-1;
        }else{
          L[[poslista]]<-c(L[[poslista]],1);
        }
      }
      prev<-epi[posprotein]
      posprotein<-posprotein+1;
    }
    L
  }
  
  shuffleAndExpandEpitope<-function(L,s=TRUE){
    ids<-1:length(L)
    if(s){
      ids<-sample(ids)
    }
    epi<-c()
    for(i in 1:length(ids)){
      epi<-c(epi,L[[ids[i]]])
    }
    epi
  }
  
  d1<-rep(NA,N)
  d0<-rep(NA,N)
  L<-collapseEpitope(epitopeAnnotation)
  for(i in 1:N){
    sepi<-shuffleAndExpandEpitope(L)
    d1[i]<-mean(entropy[which(sepi==1)],na.rm=T)
    d0[i]<-mean(entropy[which(sepi==0)],na.rm=T)
  }
  D<-mean(entropy[which(epitopeAnnotation==1)],na.rm=T)-mean(entropy[which(epitopeAnnotation==0)],na.rm=T)
  if(D>=0){
    ret<-list(Delta=D,p.value=mean(d1-d0>=D))
  }else{
    ret<-list(Delta=D,p.value=mean(d1-d0<=D))
  }
  ret
}
